# Altering the ribosome exit tunnel to improve consecutive incorporation of challenging monomers

**DOI:** 10.64898/2026.06.01.729415

**Authors:** Alexandra D. Kent, Chandrima Majumdar, Katelyn A. Fitzgerald, Alexander C. Solivan, Martina Boga de Teresa, Jessica T. Vance, Alanna Schepartz, Jamie H. D. Cate

## Abstract

Ribosomes are capable of incorporating a wide array of natural and unnatural monomers into growing polymer chains, but can be stalled by monomers with constrained or non-natural backbones. Here we evaluate whether monomer-dependent ribosome stalling can be alleviated by structure-guided mutations to 23S rRNA within the exit tunnel. Ribosomes harboring an A2062U mutation are as active as wild type (WT) ribosomes when translating non-proline sequences and up to 10-fold more active when translating sequences containing up to four consecutive proline residues. High-resolution cryo-EM structures of WT and A2062U mutant ribosomes containing a polyproline nascent chain reveal that the A2062U mutation relieves an exit tunnel constriction to better accommodate a conformationally restricted peptide chain. A2062U mutant ribosomes also improve translation of sequences containing multiple, consecutive β^2^-hydroxy acids. These results provide a mechanistic basis for engineering the ribosome for improved translation of genetically encoded polymers with novel backbones.

## Main

During translation the ribosome primarily uses a canonical set of 20 natural L-α-amino acids to generate sequence-defined polymers in the form of peptides and proteins. There has been a longstanding effort to expand the substrate scope of the ribosome to include non-canonical α-amino acids and other monomers whose backbones and side chains diverge from those found in nature^1–7^. While efforts to incorporate non-canonical α-amino acids into peptides and proteins has been highly successful^5,8^, efforts to incorporate monomers with divergent backbones and/or nucleophiles (“non-α-amino acid” monomers) have yielded varying results^5–7,9–11^. The success of these efforts have varied with monomer identity and reactivity, as well as how effectively the corresponding acylated tRNA engages with translation factors and is accommodated within the ribosome peptidyl transferase center (PTC)^9,12–16^. Thus efforts to introduce non-α-amino acids into peptides and proteins remain an ongoing goal, as polymers containing monomers with backbone extensions, such as β^2^- and β^3^- amino and hydroxy acids, can display highly desirable properties including proteolytic stability, unique architectures, and recognition of otherwise recalcitrant targets^10,17–20^. Strategies to improve the incorporation of β^2^-and β^3^ backbone monomers into peptides *in vitro* include efforts to improve EF-Tu engagement with charged tRNAs, improve reactivity of monomers within the peptidyl transferase center (PTC), and the addition of factors such as EF-P and ABC-F proteins^21–24^. Despite these advances, the introduction of multiple consecutive β^2^- and β^3^- amino and hydroxy acid backbones into peptides by the ribosome has generally suffered from low yield and scope and remains a key challenge^21,22^.

A second approach has been to modify the ribosome itself. A number of variant *Escherichia coli* (*E. coli*) ribosomes with combinatorial mutations in the vicinity of the PTC have been reported to improve incorporation of D-amino acids, *N*-Me-α-amino acids, β^3^-amino acids, and even dipeptide analogs^5,6,25–28^. Importantly, the mutant ribosomes described thus far have been either less active than wild-type (WT) *E. coli* ribosomes, misfolded when isolated and characterized *in vitro*^29^, or their function confounded by challenges with orthogonal translation systems analyzed in the presence of WT ribosomes^30,31^. It was recently reported that ribosomes from *Alteromona macleodii* (*A. macleodii*) could incorporate certain D-α- and D-β-amino acids into peptides more efficiently than *E. coli* ribosomes^32^. Notably, the rRNA sequence of the *A. macleodii* PTC is identical to that of *E. coli*, highlighting the need for deeper understanding of ribosome structure and function to improve non-natural backbones into proteins. Thus, a strategy for engineering a mutant ribosome with enhanced activity for “non-α-amino acids” while maintaining robust peptidyl transferase activity remains to be established.

This work describes a structure-guided approach to this goal. Previous structures of the bacterial ribosome that contain a well-resolved nascent peptide revealed an intricate network of hydrogen bond interactions between the backbone of the nascent chain and a small set of 23S rRNA nucleotides within the ribosomal exit tunnel. These interactions include those between two 23S rRNA nucleotides–A2062 and U2506–in a well-characterized constriction of the exit tunnel and the residue of the nascent chain located two amino acids upstream of the P-site tRNA within the PTC^33^ (**Fig. 1a**). When this residue is a conformationally constrained amino acid such as proline, modeling suggests an induced clash between A2062 and the proline side chain (PDB ID: 8CVL)^33^. Nucleotide variation at A2062 occurs naturally and mutations at this position arise spontaneously in strains exposed to macrolide antibiotics to confer a resistance phenotype^34–36^. We therefore hypothesized that mutation of A2062 to a smaller pyrimidine (U or C) would fully or partially release the constriction in the exit tunnel and improve translation of one or more conformationally constrained amino acids, including those with unnatural backbones. Indeed, here we demonstrate that a mutant ribosome with U in place of A at position 2062 demonstrates improved ability to introduce multiple proline residues, and multiple β^2^-hydroxy acids, into peptides with no effect on the translation of canonical α-amino acids.

**Figure 1:**
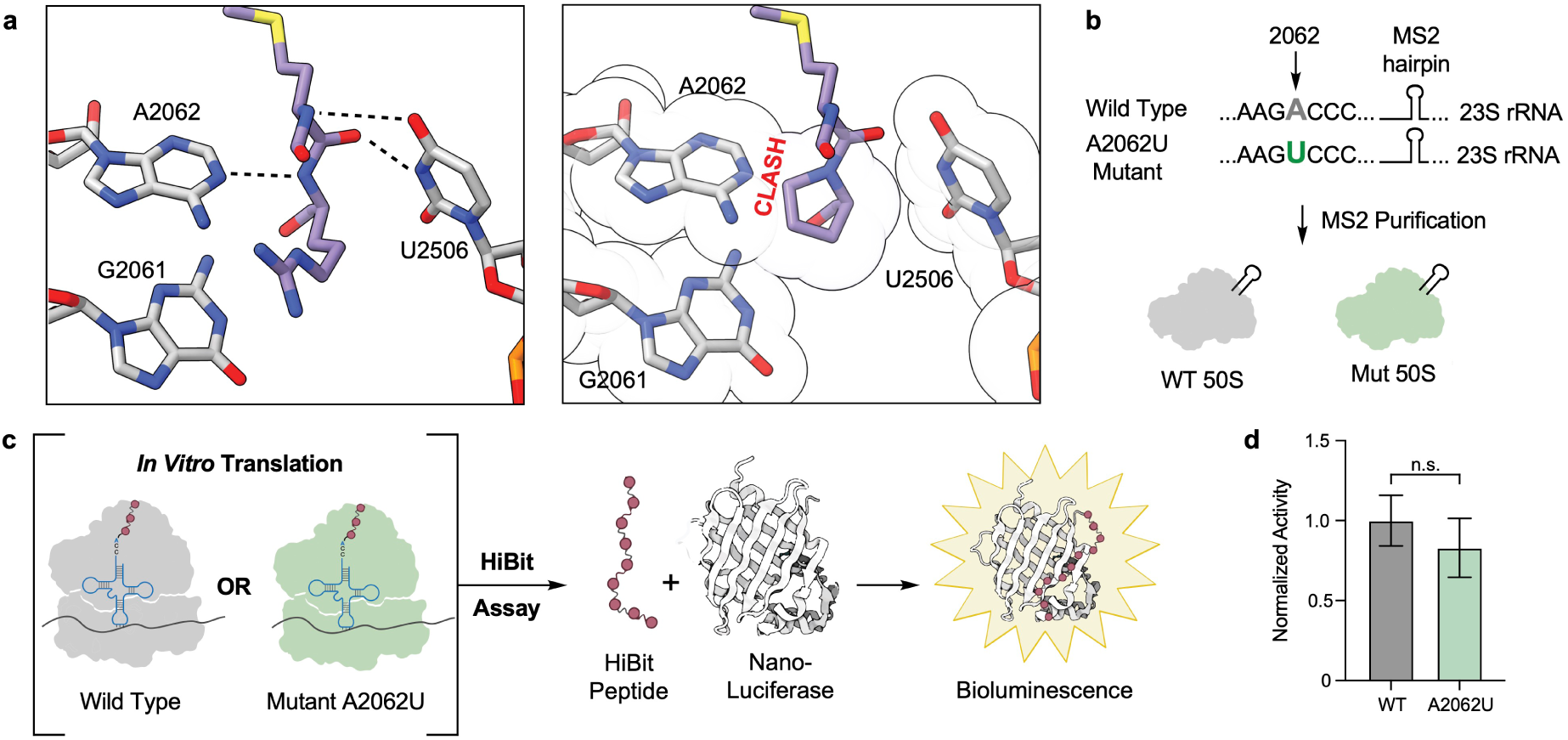
Design and rationale for an *E. coli* ribosome with a mutation of the rRNA in the exit tunnel. **a,** On the left, a 3D model of a peptide (purple) in the exit tunnel of the ribosome (grey), and on the right, a space filling model (transparent overlay) with proposed clash between a proline and A2062 (adapted from reference ^33^ PDB ID: 8CVL) **b,** Schematic of ribosomes with an MS2 hairpin added to the 23S rRNA of wild type (WT: grey) and mutant (A2062U: green) MS2-tagged ribosomes. **c,** Either WT or mutant ribosomes are used in an *in vitro* translation assay to generate the short HiBit peptide that complements the LgBit of Nanoluciferase resulting in bioluminescence. **d,** The bioluminescence is used as a readout for ribosome activity as shown in the bar graph where the y-axis is normalized activity. Error bars represent three independent replicates and the p-value was calculated to be not significant.

High-resolution cryo-EM structures of the WT and mutant *E. coli* ribosomes with a polyproline peptide in the exit tunnel provide a clear structural rationale for this improved activity. This work validates a targeted, structural approach to ribosome engineering for improved cotranslational incorporation of non-natural monomers.

## Results

### Generating A2062U mutant ribosome and assessing activity

We first sought to assess how the activity of a mutant *E. coli* ribosome containing an A2062U mutation would compare to that of the WT *E. coli* ribosome. To determine relative activities *in vitro*, we transformed *E. coli* with a plasmid whose rRNA operon encoded either the WT 23S rRNA sequence or the mutant A2062U sequence; in both cases the expressed 23S rRNA sequence contained an MS2 RNA hairpin to facilitate subunit purification and reassembly into 70S ribosomes (**Fig. 1b**). We tested the relative *in vitro* activities of the two purified ribosomes by adding them, in equal concentration, to IVTT (*in vitro* transcription translation) reactions prepared using the PURExpress® Δ Ribosome Kit and a DNA template encoding the HiBit peptide (**Supplementary Table S4**). When expressed *in vitro*, the HiBit peptide effectively complements a truncated derivative of NanoLuciferase known as LgBit to generate a bioluminescence signal (**Fig. 1c**)^13,37^. This supplementation experiment revealed that WT and A2062U ribosomes possess comparable activities (**Fig. 1d**), providing a strong starting point for evaluating the incorporation of both α- and non-α-amino, and hydroxy acid monomers.

### Ribosomes with mutation A2062U improve incorporation of sequential prolines

It is well known that polypeptide sequences rich in proline translate slowly and can stall, especially in the absence of the translation factor EF-P^38,39^. Slow translation could result from decreases in the rate of bond formation^40^. But stalling requires multiple consecutive prolines, and may also result from the compounding effects of decreased conformational flexibility and recently identified steric clashes with residue A2062 within the exit tunnel^33^. To assess the relative ability of WT or A2062 mutant ribosomes to translate through polyproline sequences, we supplemented IVTT reactions prepared using the PURExpress® Δ Ribosome Kit with equal concentrations of each ribosome and a DNA template encoding sequences with between 2 and 4 adjacent proline residues. Each translated sequence terminated in a C-terminal FLAG tag to aid subsequent analysis (**Fig. 2a,b**). IVTT reactions were incubated for four hours at 37 °C and the resulting FLAG-tagged peptides isolated using an anti-FLAG antibody coupled to magnetic beads for high resolution LC-MS analysis (**Fig. 2c-n**). The relative yield of each proline-containing peptide in reactions containing WT or mutant ribosomes was calculated from the ratio of the integrated extracted ion chromatograms of the product +5 ion.

**Figure 2:**
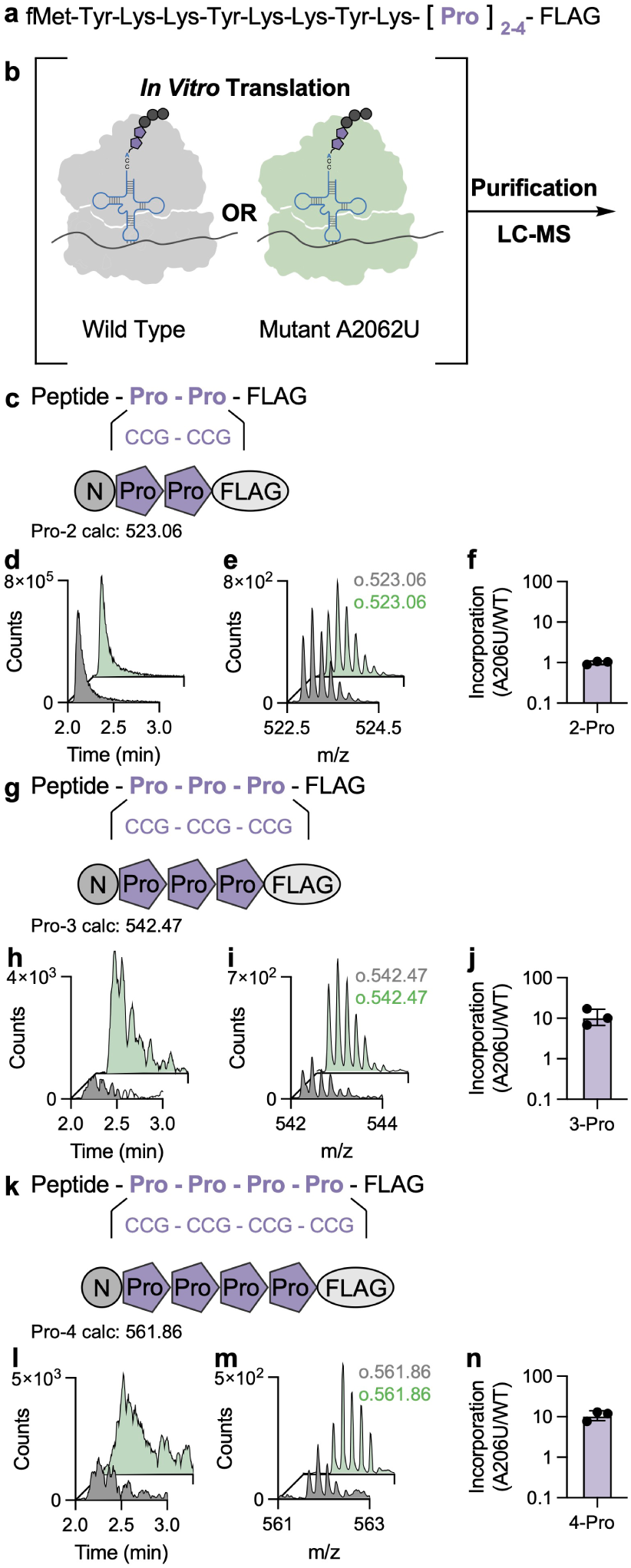
Ribosomes with an A2062U mutation incorporate polyprolines more efficiently than WT ribosomes. **a,** Sequence of the peptides produced in IVTT experiments, which contain between 2 and 4 sequential proline residues and a C-terminal FLAG-tag to aid purification and analysis. **b,** Either WT or A2062U mutant ribosomes were used to supplement IVTT reactions programmed to synthesize the peptide shown in (**a**). The peptides, purified using magnetic beads with an anti-FLAG antibody, were analyzed using LC-MS. **c,** The peptide product containing two consecutive proline residues followed by a FLAG-tag (Pro-2). The mass shown (calc.) represents that calculated m/z for the +5 ion. **d**, Shown are the extracted ion chromatograms (EICs) of Pro-2 peptide generated during IVT using either the WT (grey) and A2062U mutant (green) ribosomes. **e,** The m/z spectra of the detected +5 ion of the Pro-2 peptide. (o. is the observed m/z for the +5 ion). **f,** Relative activity of WT and A2062U ribosomes in IVTT experiments programmed to produce the Pro-2 peptide production calculated as the fold change of A2062U/WT. **g-j,** A peptide with three consecutive prolines was analyzed using LC-MS as described in **c- f. k-n,** A peptide with four consecutive prolines was analyzed using LC-MS as described in **c-f.** Error bars are for three independent replicates.

These experiments revealed that the A2062U mutation affects polyproline translation in a manner that depends on the number of consecutive prolines. When the translated polypeptide contains two adjacent prolines, there is no difference in yield irrespective of ribosome identity: the ratio of ion counts for the 2-proline containing peptide translated by WT and A2062U ribosomes is 1 (**Fig. 2d,e,f**). However, when the translated polypeptide contains three (**Fig. 2h,i,j**) or four (**Fig. 2l,m,n**) consecutive prolines, we detect a roughly 10-fold increase in yield when the IVTT reaction is supplemented with the A2062U mutant ribosome. These results are consistent with full or partial relief of the proposed clash between A2062 and the proline pyrrolidine ring at the -2 position of the nascent polypeptide chain (**Fig. 1a**)^33^. We also generated and tested mutant ribosomes with an A2062C mutation, which hampered the incorporation of polyproline sequences relative to WT ribosomes (**Extended Data Fig. 1**).

EF-P is a highly conserved translation factor found in virtually all bacteria, and is a key factor in promoting polyproline incorporation^41^. Ribosome profiling experiments performed with Δ*efp* strains that lack EF-P have identified discrete ribosome populations that are stalled genome-wide at PPP and PPX/XPP motifs^42^. Consecutive proline residues located at the C-terminus of the nascent chain are predicted to disturb peptidyl-tRNA positioning in the ribosome PTC, thereby inhibiting subsequent peptide bond formation. EF-P binds to ribosomes stalled at polyproline sequences, bridging the large and small subunits. It functions by interacting with the P-site tRNA, specifically the 3′-CCA-end, where a post-translationally modified lysine of EF-P forms stabilizing contacts and repositions the tRNA and nascent chain into a more favorable position^43^.

To evaluate whether the effects of EF-P are influenced by the A2062U mutation, we performed an analogous set of IVTT reactions that included 5 µM EF-P. In the presence of EF-P, the yields of all proline-containing products increased by 10-100 fold, but there was no additional improvement in reactions containing A2062U mutant ribosome (**Extended Data Fig. 2**). In fact, WT ribosomes are 2-3 fold more efficient than A2062U ribosomes in the presence of EF-P suggesting that the mutation may have a slight impact on the mechanism of EF-P. Additionally, EF-P also increases yields of proline-containing peptides by A2062C ribosomes, but exhibits the same rescue defect relative to WT ribosomes (**Extended Data Fig. 3**). Overall, we determined that the A2062U mutant ribosome significantly improves the incorporation of sequential proline residues without need for EF-P supplementation.

### Cryo-EM of WT and A2062U mutant ribosomes provides a structural basis for improved polyproline incorporation

To understand the structural basis for improved polyproline translation by A2062U mutant ribosomes, we devised a stalling strategy to isolate ribosomes containing a polyproline nascent chain for analysis by cryo-EM. This strategy installed a tryptophan codon following codons encoding three consecutive proline residues^44^. This design forces the ribosome to stall on the sequence of interest during an *in vitro* translation reaction that lacks tryptophan, allowing purification of fully assembled 70S ribosomes stalled with a proline-rich nascent chain within the exit tunnel (**Fig. 3a**). To test this strategy, we generated a DNA sequence with three consecutive proline codons similar to the constructs reported in **Fig. 2**, but with a tryptophan codon directly following the prolines and coding for a C-terminal FLAG-tag (**Fig. 3b**). Using a PURE system where we added a custom amino acid mixture containing tryptophan, we see production of the encoded peptide as the +5 ion in the m/z spectra with counts comparable to constructs in **Fig. 2** containing three prolines. However, when we remove tryptophan from the amino acid mixture, we see no m/z corresponding to production of the full length peptide (**Fig. 3c**).

**Figure 3:**
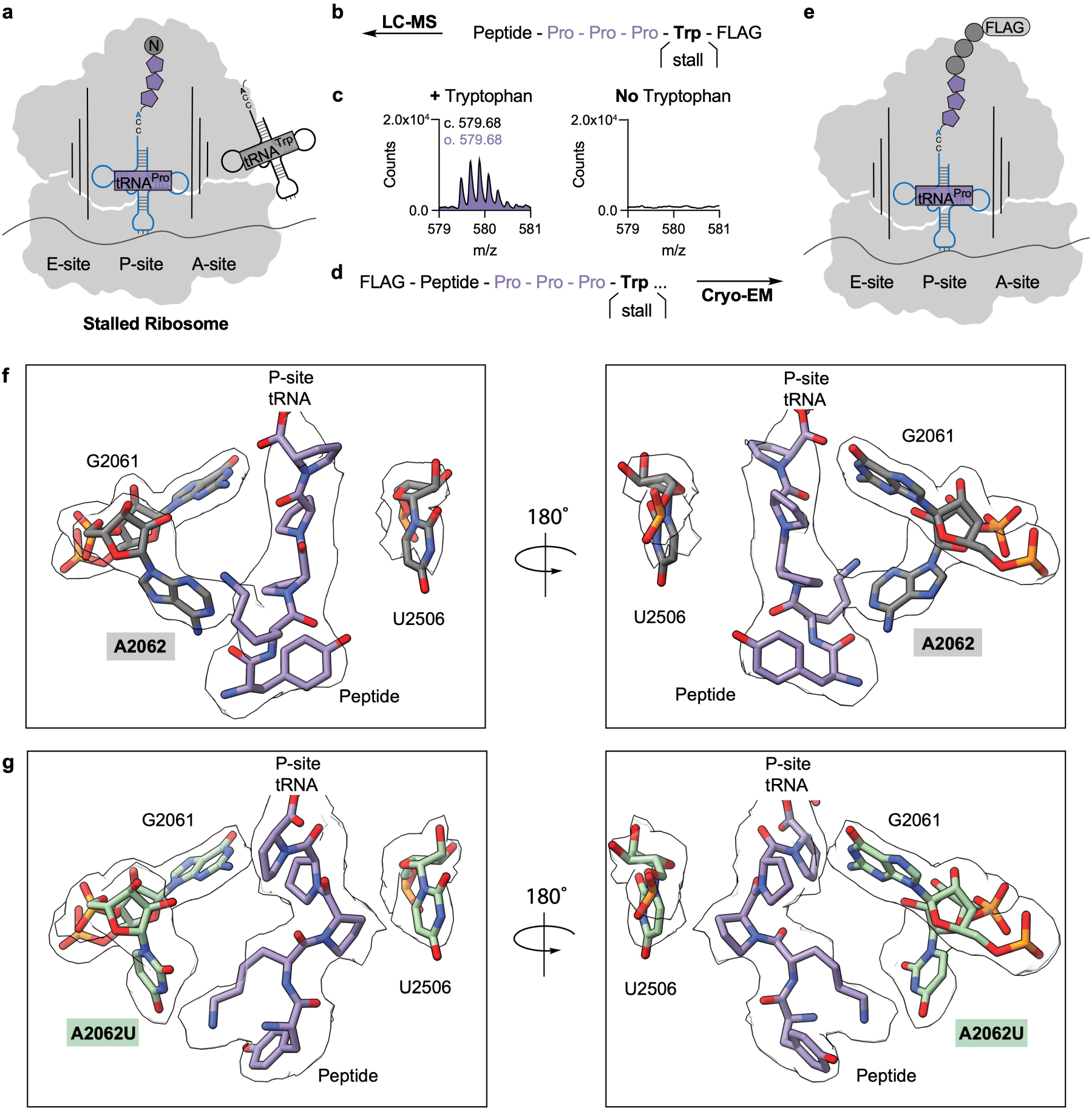
Cryo-EM of WT and A2062U mutant ribosomes with a polyproline sequence in the exit tunnel. **a,** During IVT a ribosome can incorporate three consecutive prolines, but when it encounters a tryptophan codon in the absence of tryptophan, the ribosome stalls at the third proline. **b,** Peptides that incorporate three consecutive prolines and stall on tryptophan with the FLAG-tag on the C-terminus were used for LC-MS analysis. **c,** LC-MS analysis of IVT reactions producing the peptide shown in **b** the m/z spectra of the +5 ion is shown either in the presence of tryptophan (peptide present) or absence of tryptophan (no peptide synthesized, ribosomes stalled). (c.) is the calculated m/z of the +5 ion and (o.) is the observed m/z of the +5 ion. **d**, Peptides that incorporate three consecutive prolines and stall on tryptophan with the FLAG-tag on the N-terminus were used for Cryo-EM. **e,** Ribosome stalled on the polyproline sequence with an N-terminal FLAG-tag extending out of the exit tunnel of the ribosome, allowing for purification of stalled ribosomes. **f**, Cryo-EM of WT ribosomes (grey) with a tRNA (blue) attached to a peptide containing three prolines (purple) stalled in the exit tunnel of the ribosome and a 180° rotated view. **g,** Cryo-EM of A2062U mutant ribosomes (green) with a tRNA (blue) attached to a peptide containing three prolines (purple) stalled in the exit tunnel of the ribosome and a 180° rotated view. Note: the displayed maps were supersampled for smoothness.

After confirming that tryptophan depletion establishes a viable stalling strategy, we redesigned the DNA sequence to incorporate the FLAG-tag on the N-terminus to allow for ribosome isolation (**Fig. 3d,e**)^45^. This allowed us to purify either WT or A2062U mutant 70S ribosomes stalled on a P-site tRNA^Pro^ linked to the nascent chain containing the polyproline sequence. We used single particle cryo-EM to obtain structures of the WT (1.88 Å global resolution) and mutant A2062U (2.49 Å global resolution) (**Supplementary Figs. S1-4** and **Supplementary Tables S1,2**) ribosomes allowing us to model the polyproline nascent chain in the exit tunnel and interactions with specific rRNA bases including A2062 or U2062. Resolution was improved through classification to remove particles with an occupied A-site tRNA and further classification on the nascent chain. This process allowed us to see clear density for the nascent chain in both structures.

The density for the polyproline sequence in the WT ribosomes indicates that it adopts an *trans*-like extended conformation similar to previously reported structures (**Fig. 3f**) ^43^. The structure shows clear density for rRNA base A2062 and comparison of this structure with a structure of the ribosome with a non-proline containing nascent chain shows that A2062 has shifted away from the PTC to accommodate the prolines (**Extended Data Fig. 4**). In the mutant A2062U ribosome structure, the density for nucleotide A2062U indicates that it has moved even farther away from the nascent chain, creating more space to accommodate the more favored conformation adopted by the peptide in the exit tunnel (**Fig. 3g**) ^46,47^. The density of the nascent chain suggests that the polyproline sequence begins to adopt a conformation consistent with the beginning of a polyproline helix (**Fig. 3g**) with *ω* dihedral angles between the proline residues consistent with *trans*-like peptide bonds. Other models for the nascent chain, with the prolines adopting either all *trans*- or all *cis* geometry, do not fit the observed cryo-EM density (**Extended Data Fig. 5**).

### Ribosomes with mutation A2062U improve incorporation of sequential β^2^-hydroxy acids

The above results imply that A2062U mutant ribosomes improve the incorporation of multiple consecutive proline residues by shifting nucleotide 2062 away from the growing nascent chain (**Fig. 3g**). This shift appears to allow A2062U ribosomes to better accommodate bulky consecutive pyrrolidine rings, and raises the possibility that other bulky residues would be better accommodated as well. To test this idea, we targeted monomers that translate inefficiently, specifically those with β^2^-backbones and bulky side chains^7,12,13,16^. Because of their backbone architecture, these monomers are likely unable to engage in the canonical H-bonding contacts to A2062 in the WT ribosome exit tunnel, and may experience clashes, as predicted for proline across from A2062^33^.

First we examined how well WT and A2062U mutant ribosomes support the singular incorporation of a series of non-canonical monomers with α- or β^2^-backbones and amine or hydroxyl nucleophiles. To do so, we supplemented IVTT reactions prepared using the PURExpress® Δ Ribosome Kit with equal concentrations of each ribosome, a DNA template encoding a short peptide terminating in a C-terminal FLAG tag, and acylated derivatives of a tRNA^Pyl^ variant in which the native CUA anticodon was changed to UCG (tRNA^Pyl(Ser)^). Our initial experiments focused on reactions in which tRNA^Pyl(Ser)^ was acylated with either L-α-BocLysine (α-BocK), (*R*)-β^2^-hydroxy-N^ε^-BocLysine [(*R*)-β^2^-OH-BocK], or either (*R*)-or (*S*)-β^2^-NH_2_-Phenylalanine (β^2^-NH_2_-Phe). tRNA^Pyl(Ser)^ was acylated with α-BocK and (*R*)-β^2^-OH-BocK using *M. alvi* PylRS and with (*R*)-or (*S*)-β^2^-NH_2_-Phe using the corresponding cyanomethylesters (CME) and flexizyme (**Extended Data Fig. 6**)^7,12,48^. IVTT reactions containing equal concentrations of WT and A2062U mutant ribosomes and equal concentrations of acylated tRNA^Pyl(Ser)^ were incubated for four hours at 37 °C and purified as described above before analysis by tRNA LC-MS (**Fig. 4a**)^49^.

**Figure 4:**
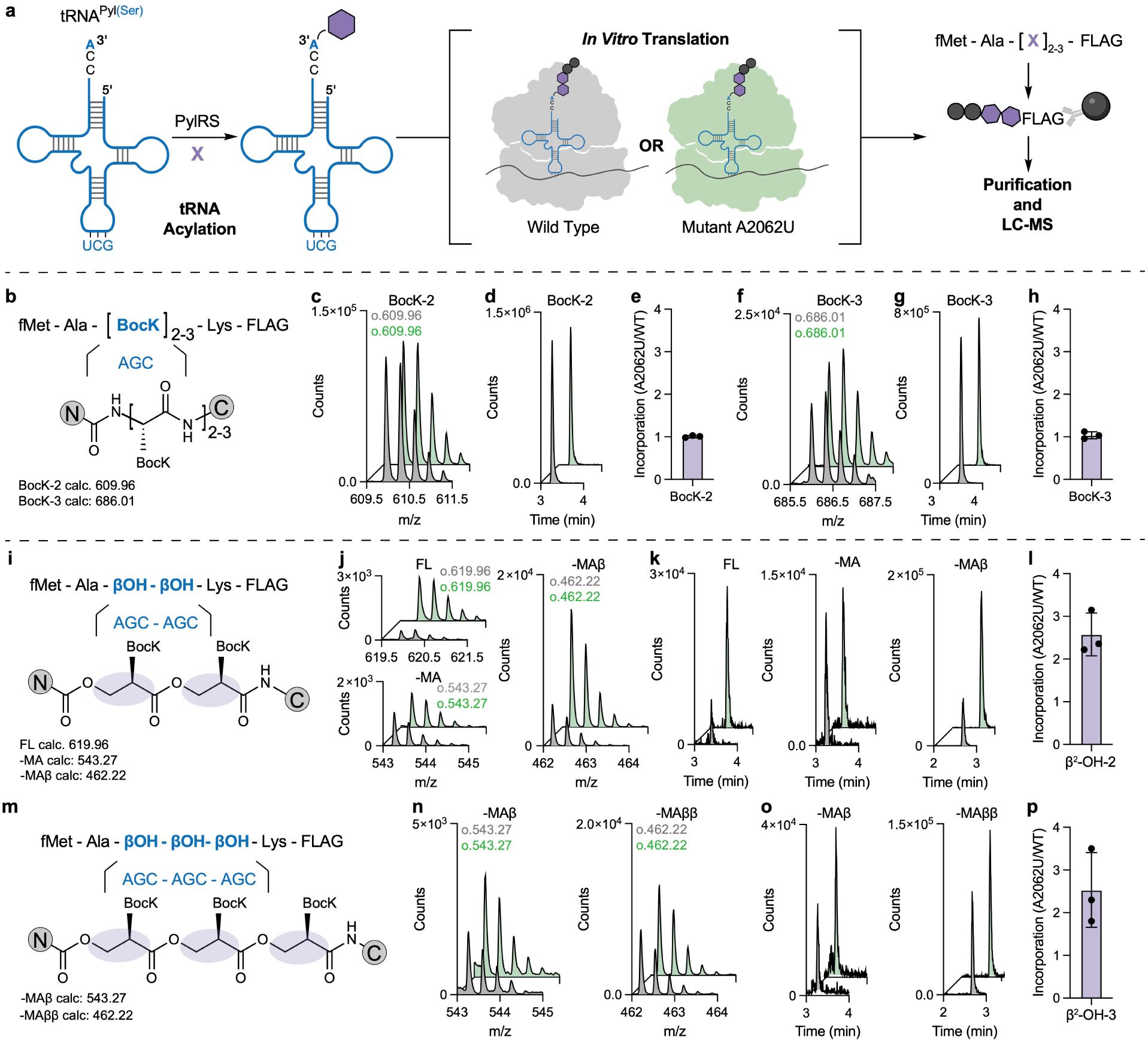
The A2062U mutant ribosome incorporates consecutive β^2^-hydroxy acids into peptides more efficiently than WT ribosomes. **a,** Reaction scheme. The *M. alvi* pyrrolysine tRNA (tRNA^Pyl^) was modified to contain the anticodon that encodes Ser (tRNA^Pyl(Ser)^) and acylated enzymatically with either α-BocK or (*R*)-β^2^-OH-BocK. These acylated tRNAs were used to supplement IVTT reactions containing either WT or A2062U mutant ribosomes and programmed to translate short peptides containing either 2 or 3 consecutive non-canonical monomers followed by a FLAG tag; the resulting polypeptide products were purified and analyzed by LC-MS. **b,** Sequence and composition of the IVTT products containing two (BocK-2) or three (BocK-3) consecutive α-BocK monomers. calc. represents the calculated m/z of the corresponding +3 ion. **c,** The m/z spectra of the +3 ion of the BocK-2 peptide product generated by either WT (grey) or 2062U mutant (green) ribosomes. The (o.) represents the observed m/z of the +3 ion. **d,** Extracted ion chromatograms (EICs) of the Boc-2 peptide product. **e,** Relative yield of the BocK-2 product calculated as the fold change in ion counts of the +3 ion in reactions containing A2062U or WT ribosomes. **f,** The m/z spectra of the +3 ion of the BocK-3 peptide product. **g,** EICs of the BocK-3 peptide product. **h,** Relative yield of the BocK-3 product calculated as the fold change in ion counts of the +3 ion in reactions containing A2062U or WT ribosomes **i,** Sequence and composition of the IVTT product containing two consecutive (*R*)-β^2^-OH-BocK monomers. **j,** The m/z spectra of the +3 ion and **k**, the EICs of the full-length product containing two consecutive (*R*)-β^2^-OH-BocK monomers and the -MA and -MAβ hydrolysis products. FL represents the full length peptide, -MA represents the C-terminal portion of the peptide following hydrolysis of the N-terminal fMet-Ala portion of the peptide, -MAβ represents the C-terminal portion of the peptide following hydrolysis of the N-terminal fMet-Ala-βOH portion of the peptide **l,** Relative peptide production calculated from all products as the fold change of A2062U/WT. **m-p,** Peptide product generated from incorporation of three consecutive (*R*)-β^2^-OH-BocK monomers. Error bars are representative of three independent replicates.

These experiments revealed that A2062U mutant ribosomes provided no benefit over WT ribosomes when incorporating a single copy of α-BocK, (*R*)-β^2^-OH-BocK, or (*S*)-β^2^-NH_2_-Phe, and hindered the incorporation of a single copy of (*R*)-β^2^-NH_2_-Phe (**Extended Data Fig. 7**). Overall, we determined that the β^2^-NH_2_-Phe monomers were incorporated into peptides by WT ribosomes approximately 100-times less efficiently than was α-BocK and 10-fold less efficiently than (*R*)-β^2^-OH-BocK. In the case of (*R*)-β^2^-OH-BocK, the products were detected as a mixture of full-length (FL) peptide and a hydrolysis product lacking the *N*-terminal two residues (-MA).

We did not detect any product when WT or A2062 mutant ribosomes were used to incorporate multiple copies of either β^2^-NH_2_-Phe enantiomer, presumably due to signals below our detection limit. Thus, we moved forward with experiments to evaluate multiple incorporation of α-BocK and (*R*)-β^2^-OH-BocK, one with an α-backbone (α-BocK) and one with an extended backbone [(*R*)-β^2^-OH-BocK].

To evaluate how well WT and A2062U mutant ribosomes support the incorporation of consecutive copies of α-BocK and (*R*)-β^2^-OH-BocK, we designed a DNA template which when transcribed would contain 2 or 3 adjacent serine codons followed by an encoded FLAG tag to enable detection of the peptide product using LC-MS (**Fig. 4a**). When translation reactions were programmed to incorporate two (**Fig. 4c-e**) or three (**Fig. 4f-h**) consecutive copies of α-BocK, the A2062U mutant ribosome was equivalent to WT ribosomes in yield. We then moved on to evaluate translation reactions programmed to incorporate two or three consecutive copies of (*R*)-β^2^-OH-BocK to establish a short polyester segment within a longer polyamide. We began with a DNA template encoding two consecutive (*R*)-β^2^-OH-BocK monomers (**Fig. 4i**), and detected the expected peptide (FL), along with two apparent hydrolysis products corresponding to hydrolysis of one or both ester bonds during the low pH peptide purification procedure (**Fig. 4j,k**). Control experiments confirmed that these products resulted from neither serine mis-incorporation nor non-AUG initiation (**Extended Data Figs. 8,9**). We summed the integrated EICs for each product to determine overall relative incorporation efficiency and determined that the mutant A2062U ribosome incorporates two consecutive (*R*)-β^2^-OH-BocK monomers approximately 2.5-fold more efficiently than WT ribosomes (**Fig. 4l**). Finally, we incorporated three consecutive (*R*)-β^2^-OH-BocK monomers and determined that the mutant A2062U ribosome also improves incorporation of three β^2^-monomers by approximately 2.5-fold (**Fig. 4m-p**). These data demonstrate that mutation A2062U in the exit tunnel of the ribosome improves consecutive incorporations of (*R*)-β^2^-OH-BocK monomers to form short polyesters.

## Discussion

The challenges associated with engineering the ribosome for new functions are considerably more complex than engineering the chemistry of stand-alone enzymes, which has witnessed remarkable successes over the last decade^50^. First, the sequence space associated with even the most compact ribosomes is vast: the large subunit rRNA of the *E. coli* ribosome contains close to 3000 bases, and thus combinatorial mutagenesis remains beyond the reach of even continuous evolution schemes. Moreover, a large fraction of the residues located within the PTC are universally conserved and are involved in complex allosteric networks that are only partially understood^32,51,52^. Unlike many stand-alone enzymes, translation requires multiple processes, involving numerous translation factors and regulatory mechanisms that have evolved for the distinct purpose of maintaining the accurate biosynthesis of biopolymers containing only α-amino acids^53^.

Previous attempts to engineer the ribosome for new chemistry have approached these challenges by screening libraries with mutational diversity in various segments of the PTC.^11^ Ribosomes with multiple mutations within the PTC can introduce β^3^-amino acids, dipeptides and heterocycles derived thereof, *N*-methyl-amino acids, and certain D-α and D-β-amino acids *in vitro* ^5,6,26,27,30^. However, none of these mutant ribosomes have been characterized at high-resolution to provide an understanding of the emergent function; the one ribosome for which structural information is available was largely unfolded within the mutated region when visualized by cryo-EM^29^. Furthermore, mutant ribosomes reported to improve polyproline used an orthogonal ribosome expressed in the presence of WT ribosomes^30^. Although orthogonality of these ribosomes was improved, some WT cross-reactivity still occurred in cells, and was not rigorously excluded. Furthermore, the causative mutation or mutations responsible for the reported improvement in polyproline incorporation were not identified experimentally, or compared directly with incorporation by WT ribosomes. Finally, subsequent work also showed that orthogonal mRNAs used with orthogonal ribosome systems may not be sufficient to prevent translation by WT ribosomes in the cell^31^.

Here we approach the challenges associated with ribosome evolution through the lens of structural biology. We were inspired by a close examination of how a nascent chain composed of canonical α-amino acids progresses through the ribosome beginning with peptide bond formation and subsequent elongation through the exit tunnel. Certain sequences of amino acids, such as in SecM and TnaC, or even polyproline stretches, are known to induce ribosomal stalling through various mechanisms, some of which involve the exit tunnel^43,54,55^. Previous work has shown that specific mutations to PTC nucleotides at or near the exit tunnel can relieve some translational stalls^56,57^. Further, the crystal structure of the bacterial ribosome complexed with a non-stalling peptide highlights a narrow constriction in the upper exit tunnel comprised of bases in the 23S rRNA, including A2062, that may clash with monomers having bulky backbones^33^. We tested mutation to a smaller pyrimidine base, both A2062U and A2062C, and found that A2062U significantly improved ribosome activity for translating multiple consecutive prolines and β^2^-hydroxy acids. Precedent for this mutation is also found in some bacterial strains conferring resistance to macrolide antibiotics, which function by blocking the path of the nascent peptide chain through the exit tunnel^34,36^.

Structural comparison of the WT and A2062U mutant ribosome using cryo-EM showed that, in the WT ribosome, A2062 constricts the exit tunnel and forces the growing poly-Pro sequence to adopt a more linear conformation that we modeled as *trans*-like prolines. Mutation of A2062 to U relieves this bottleneck and enables the polypeptide to adopt the beginnings of a helical structure composed of prolines with more favorable *ω* dihedral angles that bring them into closer alignment with a *trans*-proline conformation. We hypothesize that relieving this torsional strain contributes to the observed translation improvement of consecutive prolines compared to WT ribosomes. In addition to having a constrained ring structure, proline is the only natural *N*-alkyl amino acid in the genetic code. Efforts to incorporate *N*-alkyl amino acids into proteins have been challenging due to their slower kinetics of bond formation and possible cellular metabolism^40,58^. The A2062U mutation may serve as a basis for improving the incorporation of other *N*-alkyl amino acids beyond proline and enable the *in vitro* ribosomal synthesis of peptides with functionalized backbones.

In a prior study, we showed that a single β^2^-OH-BocK monomer is well accommodated by the ribosomal PTC, regardless of stereochemistry, and perfectly poised for bond formation when bound within the ribosomal A site^16^. Despite their favorable orientation, incorporation of these monomers is still limited in yield. Here, we show that increasing the space within the ribosome exit tunnel improves the incorporation of up to three *(R)*-β^2^-OH-BocK monomers into a peptide implying that this mutation may lower the barrier of a rate limiting step that follows the initial bond-forming reaction. While β^2^-hydroxy acids can be incorporated consecutively in a programmable fashion using a rationally designed mutant, our results indicate that β^2^-amino acids remain a challenging substrate for the ribosome and we were unable to detect multiple incorporations at our limit of detection. This may be as a result of the chemistry of bond formation within the ribosome. Although β^2^-amino acids and β^2^-hydroxy acids both involve the nucleophilic attack of the -NH_2_ or -OH on the carbonyl group of the P-site monomer, β^2^-amino acids may not be positioned ideally for the necessary amine deprotonation. This step may be rate limiting and highly dependent on the stereochemistry of the monomer, indicating alternative methods may still be necessary to obtain efficient production of many β^2^-amino acids containing peptides. However, in this work we have expanded the substrate scope of the ribosome to enable the *in vitro* biosynthesis of a β^2^-polyester, enabling access to sequence defined polymers having pH-tunable properties with applications in drug-delivery and new materials with foldamer properties. Taken together, the results with A2062U ribosomes demonstrate that structure guided mutations at sites distal to the PTC may be a viable strategy towards expanding the substrate scope of the ribosome.

## Data Availability

### Supporting Information

Cryo-EM data collection and model refinement statistics, supplementary figures, and supplementary tables.

### Accession Codes

Atomic coordinates have been deposited with the Protein Data Bank under accession codes 13MA and 13MB (for the WT and Mutant A2062U *E. coli* ribosome, respectively). Cryo-EM maps have been deposited with the Electron Microscopy Data Bank under the accession codes EMD-77148 (70S map for the WT *E. coli* ribosome) and EMD-77149 (70S map for the A2062U mutant *E. coli* ribosome).

## Acknowledgements

This work was supported by the NSF Center for Genetically Encoded Materials, CHE-2002182 and CHE-2503885. We thank the Cal-Cryo Center, Dr. Dan Toso, Dr. Ravindra Thakkar, and Paul Tobias for help with cryo-EM data collection and management. We thank Yuk-Cheung Chan and the Miller Lab at Yale for synthesis of the β^2^-Phe-CME monomers and members of the Schepartz Lab for providing us with PylRS. We thank Dr. Amos Nissley for insightful conversations. We thank Prof. Yury Polikanov for helpful discussions on the cryo-EM experiments and structures.

## Author Contributions

ADK and JHDC conceived the study; ADK and JHDC designed the A2062 mutant ribosomes; ADK, KAF, MBT, and JTV purified mutant and WT ribosomes. ADK and KAF designed, optimized, and analyzed IVTT experiments for proline incorporation and EF-P effects with assistance from MBT and JTV; ADK and MBT designed and verified the ribosome stalling method for Cryo-EM; ADK and CM collected cryo-EM data with assistance from MBT; CM and ADK analyzed the cryo-EM data and modeled the structures; ADK designed and analyzed β^2^-hydroxy acid IVTT experiments with tRNA acylation optimization by KAF and JTV; ACS performed experiments with β^2^-amino acid substrates; ADK, CM, JHDC, and AS wrote the manuscript with input from all authors. ⁑ These authors contributed equally. KAF and ACS contributed equally and are listed alphabetically.

## Extended Data Figures

**Extended Data Figure 1:**
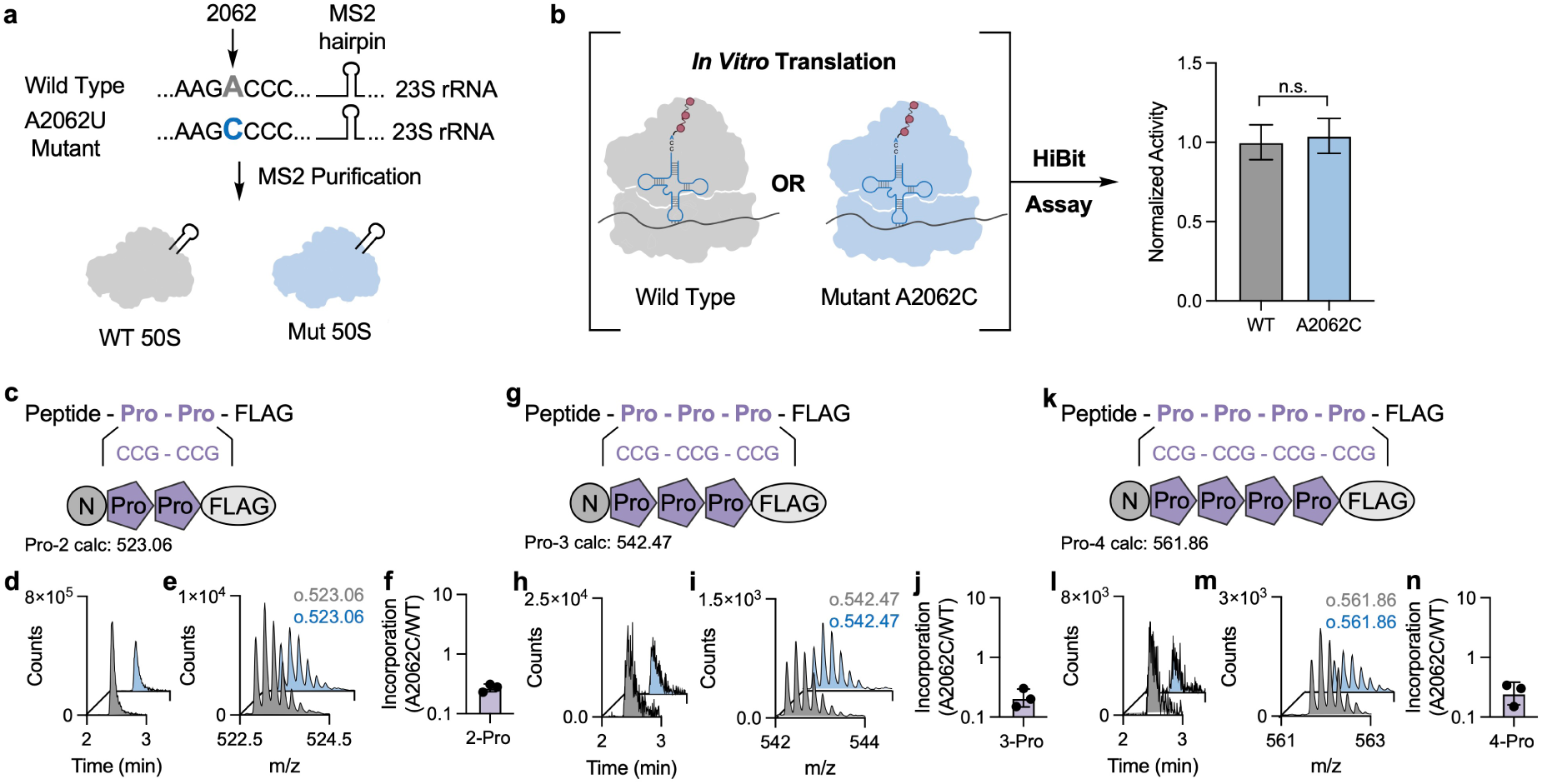
Activity of mutant A2062C ribosomes and polyproline incorporation. Mutant A2062C ribosomes incorporate consecutive polyprolines less efficiently than WT ribosomes. **a,** Mutant A2062C ribosomes (blue) were generated with an MS2 hairpin and subsequently purified. **b,** Either WT (grey) or mutant A2062C (blue) ribosomes were added to an *in vitro* translation reaction to generate a HiBit peptide. The HiBit peptide complements with the LgBit of Nanoluciferase to generate a bioluminescence signal. No difference in activity was observed between WT and mutant ribosomes. **c,** Peptide with two consecutive prolines and calculated mass for the +5 ion. **d,** Extracted ion chromatograms (EIC) of the peptide in **c** with WT ribosomes in grey and A2062C ribosomes in blue. **e,** The m/z spectra showing the +5 ion and the observed m/z values. **f,** Ratio of peptide produced (A2062C/WT). **g-j,** Peptide with three consecutive prolines, EICs, m/z spectra, and ratio of peptide produced. **k-n,** Peptide with four consecutive prolines, EICs, m/z spectra, and ratio of peptide produced. Error bars are representative of three independent experiments.

**Extended Data Figure 2:**
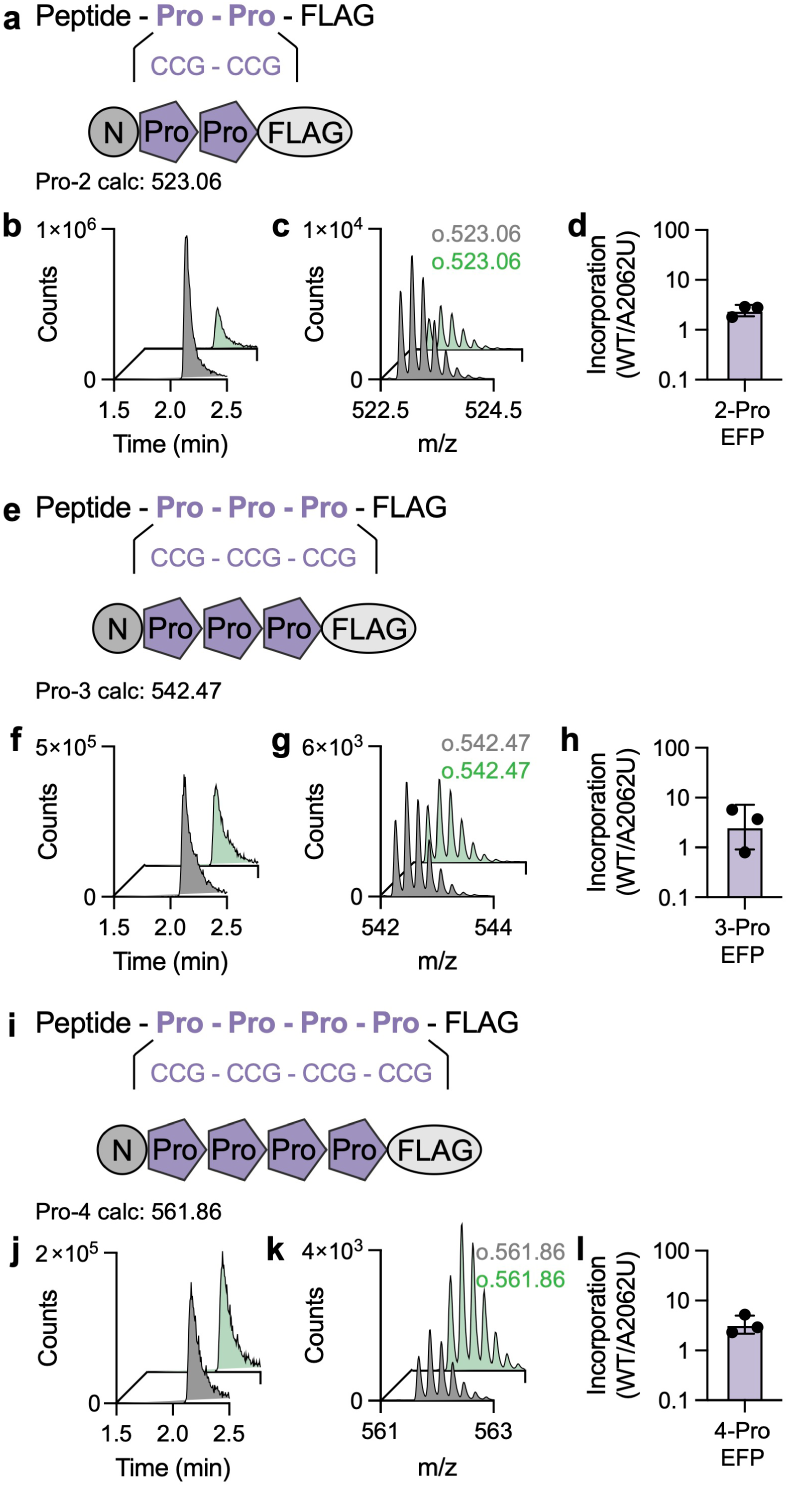
Polyproline incorporation by WT and A2062U mutant ribosomes in the presence of EF-P. Addition of EF-P significantly improves incorporation of polyproline sequences when using either WT or A2062U ribosomes. Higher incorporation is observed when WT ribosomes are used compared to A2062U mutant ribosomes. **a,** Peptide with two consecutive prolines and calculated mass for the +5 ion. **b,** Extracted ion chromatograms (EIC) of the peptide in (**a**). WT ribosomes are in grey and A2062U ribosomes are in green. **c,** The m/z spectra showing the +5 ion and the observed m/z values. **d,** Ratio of peptide produced (WT/A2062U). **e-h.** Peptide with three consecutive prolines, EICs, m/z spectra, and ratio of peptide produced. **i-l,** Peptide with four consecutive prolines, EICs, m/z spectra, and ratio of peptide produced. Error bars are representative of three independent experiments.

**Extended Data Figure 3:**
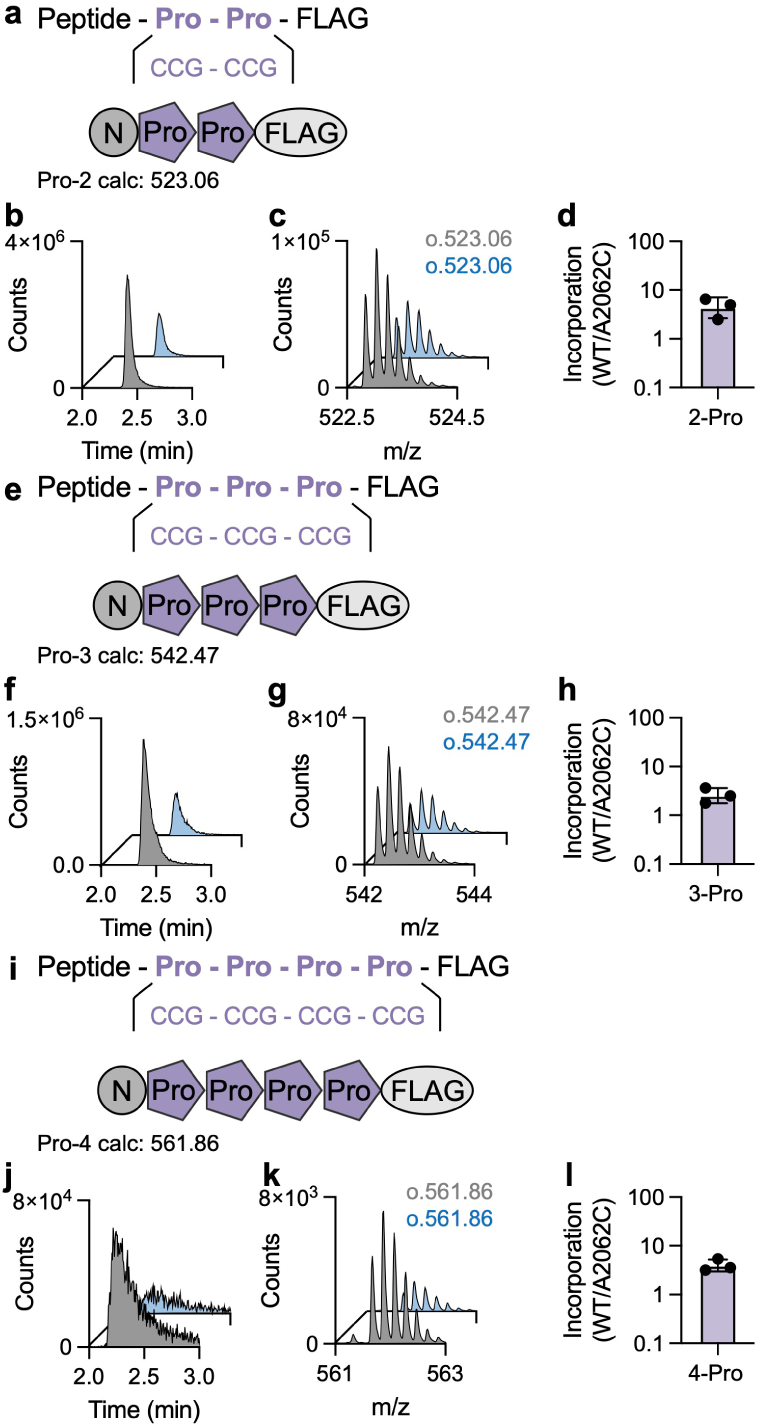
Polyproline incorporation by WT and A2062C mutant ribosomes in the presence of EF-P. Addition of EF-P significantly improves incorporation of polyproline sequences when using either WT or A2062C ribosomes. Higher incorporation is observed when WT ribosomes are used compared to A2062C mutant ribosomes. **a,** Peptide with two consecutive prolines and calculated mass for the +5 ion. **b,** Extracted ion chromatograms (EIC) of the peptide in (**a**). WT ribosomes are in grey and A2062C ribosomes are in blue. **c,** The m/z spectra showing the +5 ion and the observed m/z values. **d,** Ratio of peptide produced (WT/A2062C). **e-h,** Peptide with three consecutive prolines, EICs, m/z spectra, and ratio of peptide produced. **i-l,** Peptide with four consecutive prolines, EICs, m/z spectra, and ratio of peptide produced. Error bars are representative of three independent experiments.

**Extended Data Figure 4:**
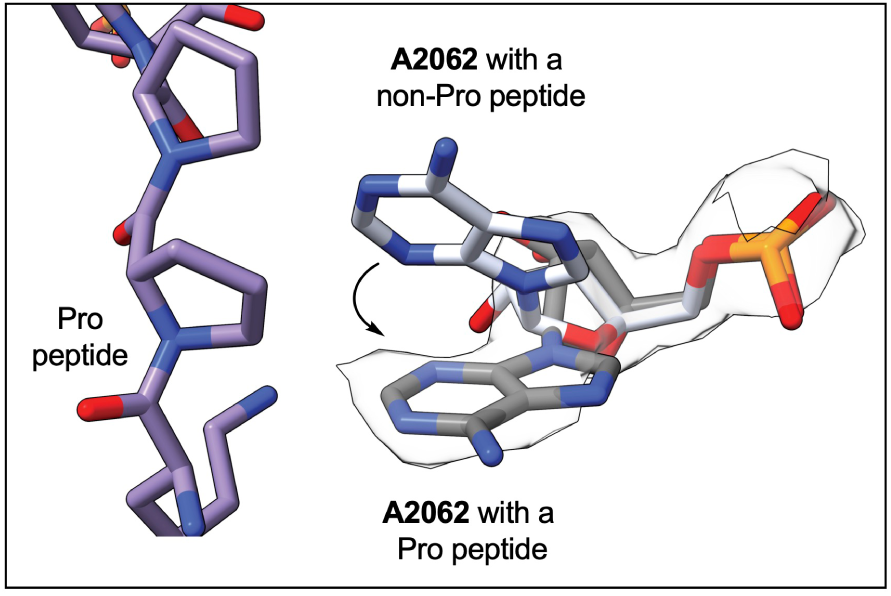
A2062 shifts position to accommodate a proline-containing peptide. A2062 modeled from a structure containing a non-Pro peptide in the exit tunnel (white) (PDB ID: 8CVL). When prolines are present in the exit tunnel (purple), A2062 (grey) shifts to accommodate the Pro-containing peptide.

**Extended Data Figure 5:**
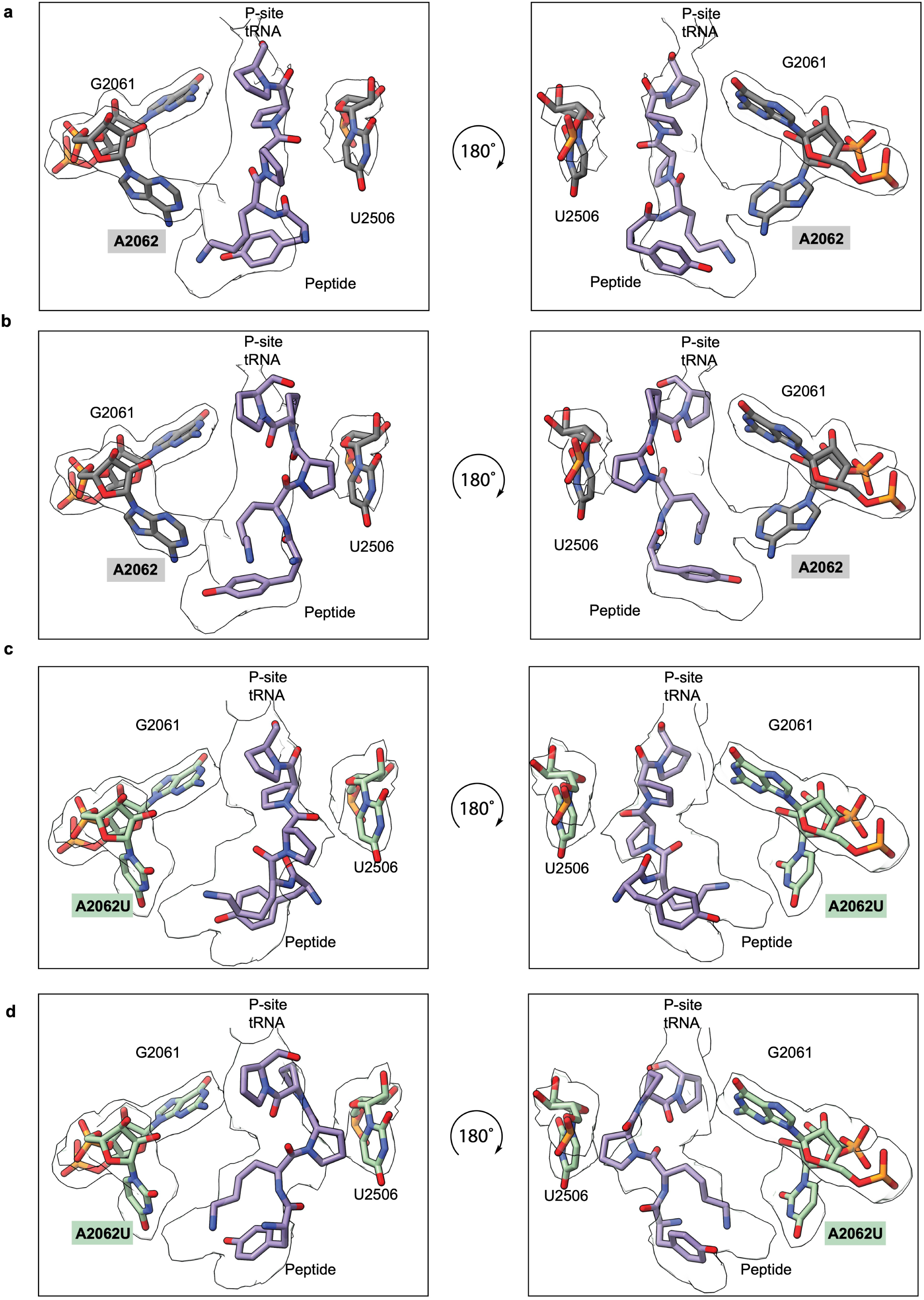
Modeling alternative conformations of the poly-proline sequence in the WT ribosome with **a,** all *trans-*proline bonds and **b,** all *cis*-proline bonds, or the A2062U mutant ribosome with **c,** all *trans-* proline bonds or **d,** all *cis-*proline bonds do not fit the observed cryo-EM density.

**Extended Data Figure 6:**
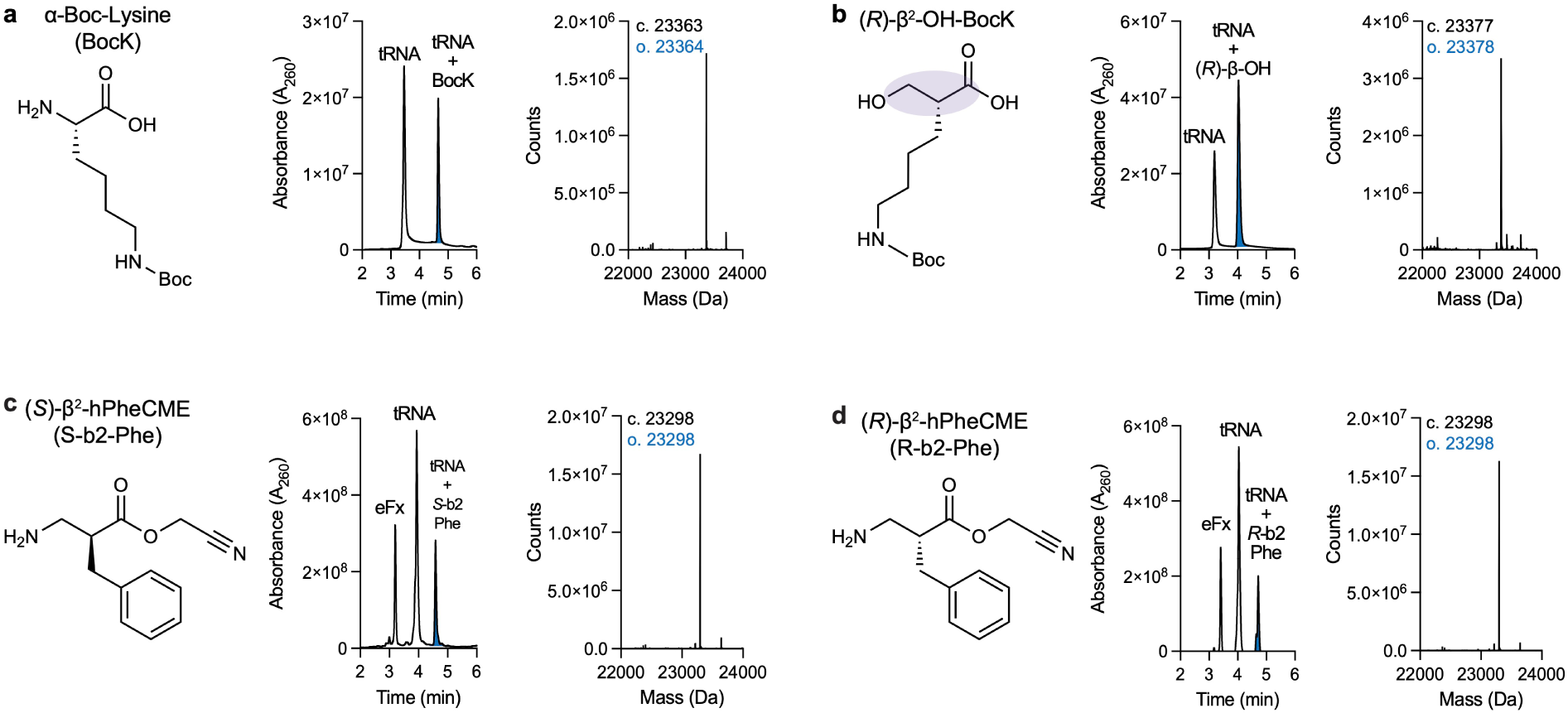
Acylation of tRNA^Pyl(Ser)^ with α-Boc-Lysine, (*R*)-ꞵ^2^-OH-BocK, (*R*)-ꞵ^2^-NH_2_-Phe, or (*S*)-ꞵ^2^-NH_2_-Phe using either PylRS or a flexizyme. **a,** The monomer α-Boc-Lysine (BocK) was acylated onto tRNA^Pyl(Ser)^ using PylRS. Shown is the LC trace measuring absorbance at 260 nm with the starting material (tRNA) and acylated product (blue, tRNA + BocK) labeled. The deconvoluted mass spectra of the acylated tRNA is shown with the calculated (c.) and observed (o.) masses shown. **b,** The monomer (*R*)-ꞵ^2^-OH-BocK R-ꞵ-OH) was acylated onto tRNA^Pyl(Ser)^ using PylRS. Shown is the LC trace measuring absorbance at 260 nm with the starting material (tRNA) and acylated product (blue, tRNA + R-ꞵ-OH) labeled. The deconvoluted mass spectra of the acylated tRNA is shown with the calculated (c.) and observed (o.) masses shown. **c,** The monomer (*S*)-ꞵ^2^-NH_2_-Phe (S-b2-Phe) was acylated onto tRNA^Pyl(Ser)^ using the flexizyme, eFx. Shown is the LC trace measuring absorbance at 260 nm with the flexizyme (eFx) starting material (tRNA) and acylated product (blue, tRNA + S-b2-Phe) labeled. The deconvoluted mass spectra of the acylated tRNA is shown with the calculated (c.) and observed (o.) masses shown. **d,** The monomer (*R*)-ꞵ^2^-NH_2_-Phe (R-b2-Phe) was acylated onto tRNA^Pyl(Ser)^ using the flexizyme, eFx. Shown is the LC trace measuring absorbance at 260 nm with the flexizyme (eFx) starting material (tRNA) and acylated product (blue, tRNA + R-b2-Phe) labeled. The deconvoluted mass spectra of the acylated tRNA is shown with the calculated (c.) and observed (o.) masses shown.

**Extended Data Figure 7:**
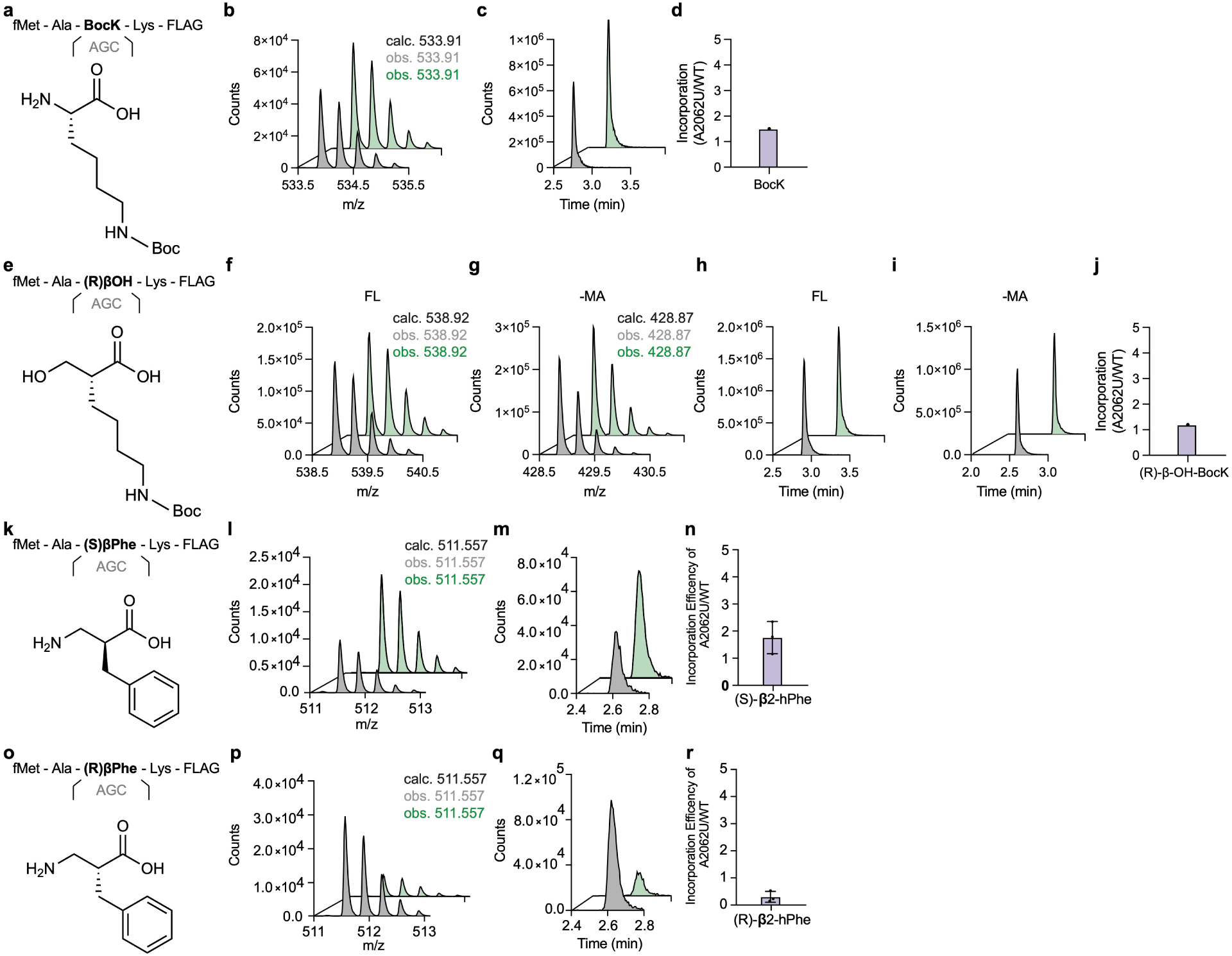
Single incorporation of unnatural monomers by WT and mutant A2062U ribosomes. Mutant ribosomes show no benefit over WT ribosomes for incorporation of one unnatural monomer. **a,** α-BocK incorporated over into a peptide over a recoded serine codon. **b,** The m/z spectra of the +3 ion with calculated and observed m/z values listed (WT: grey, A2062U: green). **c,** Extracted ion chromatograms (EICs). **d,** Ratio of peptide produced (A2062U/WT). **e,** (*R*)-ꞵ^2^-OH-BocK incorporated into a peptide over a recoded serine codon. **f,** The m/z spectra of the +3 ion of the full length (FL) peptide with calculated and observed m/z values listed. **g,** The m/z spectra of the +3 ion of the hydrolyzed peptide having lost the fMet and Ala amino acids (-MA). **h,** EICs of the FL peptide. **i,** EICs of the -MA peptide. **j,** Ratio of peptide produced calculated from the sum of both products (A2062U/WT). **k**, (*S*)-ꞵ^2^-NH_2_-Phe incorporated into a peptide over a recoded serine codon. **l,** The m/z spectra of the +3 ion with calculated and observed m/z values listed. **m,** EICs of the full length peptide. **n,** Ratio of peptide produced (A2062U/WT). Error bars are representative of three independent replicates. **o,** (*R*)-ꞵ^2^-NH_2_-Phe incorporated into a peptide over a recoded serine codon. **p,** The m/z spectra of the +3 ion with calculated and observed m/z values listed. **q,** EICs of the full length peptide. **r,** Ratio of peptide produced (A2062U/WT). Error bars are representative of three independent replicates. (*R*)-ꞵ^2^-NH_2_-Phe is incorporated less efficiently by mutant A2062U ribosomes compared to WT ribosomes.

**Extended Data Figure 8:**
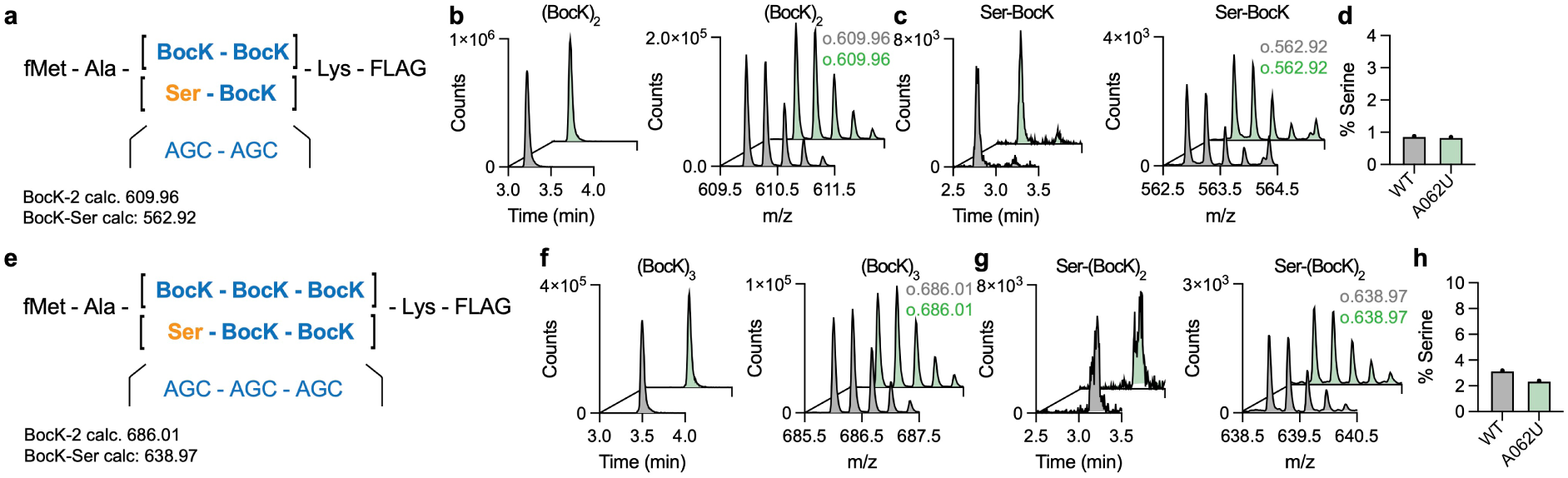
Off target incorporation of serine over the serine codon. Though serine is not added to the PURE *in vitro* translation reactions, some trace serine may be present in the reaction mixture, thus we wanted to quantify the percentage of off-target serine incorporation. **a,** In an IVT reaction lacking serine, serine codons were recoded to incorporate two consecutive α-BocK monomers over serine codons using tRNA^Pyl(Ser)^. The mass for a single and double misincorporation of serine were searched for, only single misincorporation of serine was present. **b,** Extracted ion chromatograms (EICs) and m/z spectra of the +3 ion with observed m/z values of the full length peptide with two consecutive α-BocK monomers (WT: grey, A2062U: green). **c,** EICs and m/z spectra of the +3 ion of a single serine misincorporation. **d,** Percentage of serine misincorporation seen in WT and A2062U mutant ribosomes. Similarly low percentages are present in both IVT reactions. **e,** In an IVT reaction lacking serine, serine codons were recoded to incorporate three consecutive α-BocK monomers over serine codons using tRNA^Pyl(Ser)^. The mass for a single, double, and triple misincorporation of serine were searched for, only single misincorporation of serine was present. **f,** EICs and m/z spectra of the +3 ion of the full length peptide with three consecutive α-BocK monomers. **g**, EICs and m/z spectra of the +3 ion of one serine misicorporation and two α-BocK monomers. **j,** Percentage of serine misincorporation seen in WT and A2062U mutant ribosomes. Similarly low percentages are present in both IVT reactions.

**Extended Data Figure 9:**
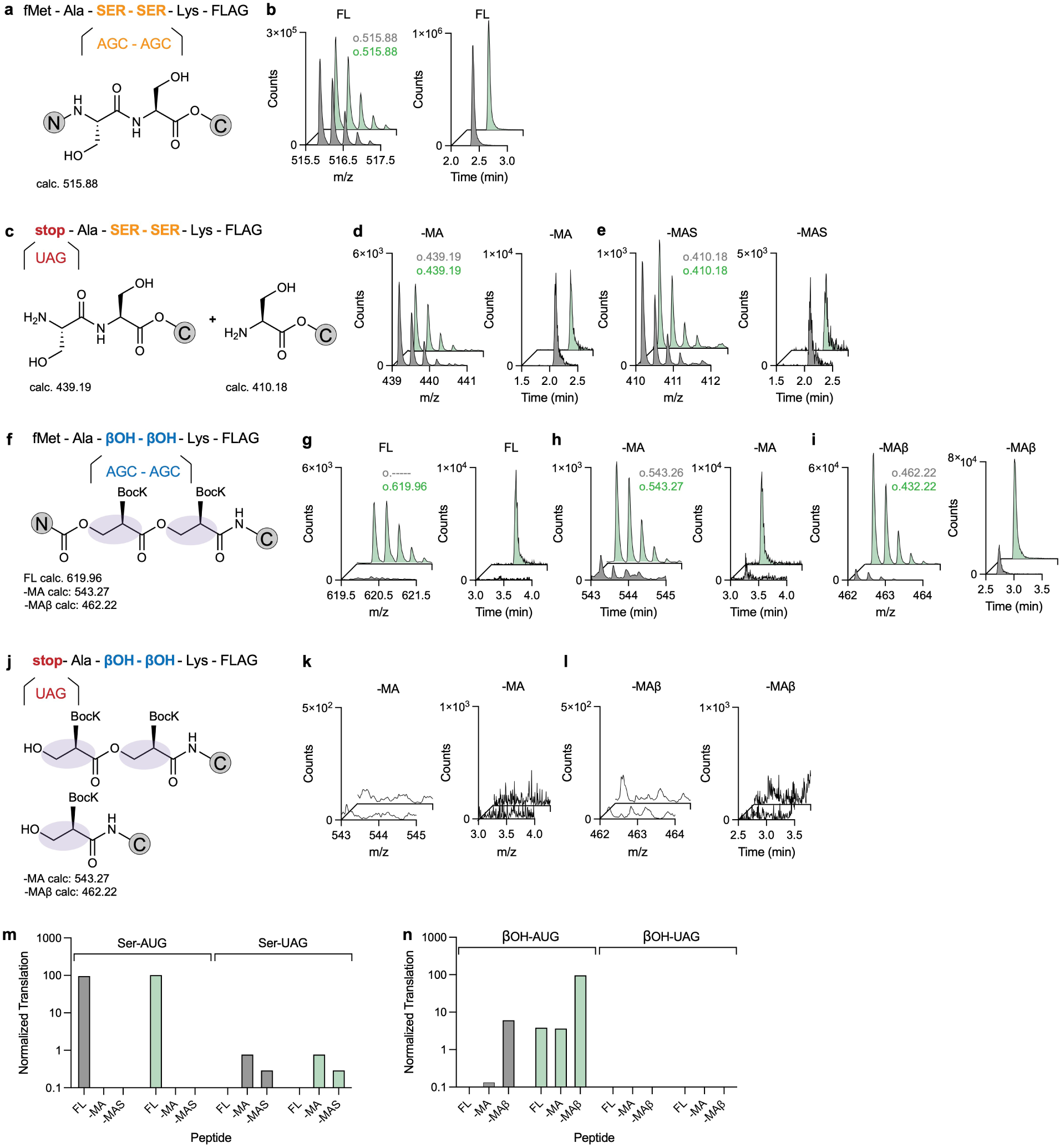
Non-start codon initiation during *in vitro* translation. A low percentage of non-canonical initiation with serine is observed when a start codon isn’t present, but there is an absence of any non-canonical initiation products with ꞵ-OH monomers. **a,** A peptide with an AUG start codon with two consecutive canonical serine incorporated over serine codons. **b,** The m/z spectra of the +3 ion and extracted ion chromatograms of the full length peptide (WT: grey, A2062U: green). **c,** A sequence for a peptide where the start codon has been replaced with a UAG stop codon to determine if alternate initiation occurs on serine codons with serine. **d,** The m/z spectra of the +3 ion and EICs of a peptide generated from initiation at the first serine codon. **e,** The m/z spectra of the +3 ion and EICs of a peptide initiating at the second serine codon. **f,** A peptide with an AUG start codon with two consecutive (*R*)-ꞵ^2^-OH-BocK incorporated over serine codons. **g,** The m/z spectra of the +3 ion and EICs of the full length peptide. **h,** The m/z spectra of the +3 ion and EICs of the hydrolyzed peptide where the fMet and Ala amino acids have been removed. **i,** The m/z spectra of the +3 ion and EICs of the hydrolyzed peptide where the fMet and Ala amino acids and one ꞵ-OH monomer have been removed. **j,** A sequence for a peptide where the start codon has been replaced with a UAG stop codon to determine if alternate initiation is occurring with ꞵ-OH monomers. **k,** The m/z spectra and EICs showing no peptide lacking the fMet and Ala amino acids has been synthesized. **l,** The m/z spectra and EICs showing no peptide lacking the fMet, Ala amino acids, and ꞵ-OH monomer has been synthesized. **m,** A bar graph showing the relative production of serine containing peptides in each experiment. **n,** A bar graph showing the relative production of ꞵ-OH containing peptides in each experiment.

## Notes

### Competing Interest Statement

A.K. and J.H.D.C. have filed a patent application related to this work. All the other authors declare no competing interests.

