## Supplementary Information for "Altering the ribosome exit tunnel to improve consecutive incorporation of challenging monomers"

### Table of Contents

**Methods**

a. *Cloning and plasmid design to generate A2062U mutant ribosomes.*
b. *Expression and purification of MS2-tagged WT and A2062U mutant ribosomes.*
c. *Quantification of activity using a bioluminescence complementation assay.*
d. *In vitro translation of peptides containing a variable number of prolines.*
e. *Purification and analysis of FLAG-tagged peptides.*
f. *Transcription of tRNA$^{Pyl(Ser)}$*
g. *Enzymatic acylation of tRNA$^{Pyl(Ser)}$ with either BocK or (R)-β$^2$-OH-BocK.*
h. *In vitro translation of peptides containing either α-BocK or β$^2$-OH-BocK.*
i. *In vitro translation of peptide to test tryptophan codon stalling strategy.*
j. *Cryo-EM complex preparation.*
k. *Cryo-EM sample preparation.*
l. *Cryo-EM data collection.*
m. *Cryo-EM image processing.*
n. *Modeling.*

**Supplementary Figures**

Figure S1: Resolution of the WT 70S ribosome with a polyproline nascent chain.
Figure S2: Resolution of the mutant A2062U 70S ribosome with a polyproline nascent chain.
Figure S3: Cryo-EM data processing workflow for the WT 70S ribosome with a polyproline nascent chain.
Figure S4: Cryo-EM data processing workflow for the mutant A2062U 70S ribosome with a polyproline nascent chain.

**Supplementary Tables**

Table 1: Data collection and processing for the WT 70S ribosome and the mutant A2062U ribosome with a polyproline nascent chain.
Table 2: Model refinement statistics for the WT 70S ribosome and the mutant A2062U ribosome with a polyproline nascent chain.

Table 3: DNA sequences.
Table 4: Peptide sequences.

**Methods**

*Cloning and plasmid design to generate A2062U mutant ribosomes.*

A modified pLK35 plasmid[1] encoding the MS2-tagged 23S rRNA was used to generate the mutant ribosomes. A point mutation at A2062 was made using the In-Fusion Cloning Kit (Takara Bio) and the corresponding primer sets (Table S3). The resulting pLK35-A2062U plasmid was cloned and sequenced to verify the presence of the A2062U mutation.

*Expression and purification of MS2-tagged WT and A2062U mutant ribosomes.*

Mutant *E. coli* 50S ribosomal subunits were expressed and crudely purified as previously described[1]. MS2-tagged 50S subunits were then purified using a maltose binding protein (MBP)–MS2 fusion protein, which was purified as previously described[2]. A 5 mL MBP Trap column (Cytiva) at 4°C was washed with five column volumes (CV) of MS2-150 buffer (20 mM HEPES pH 7.5, 150 mM KCl, 1 mM EDTA, 2 mM 2-mercaptoethanol) and then 10 mg of MBP–MS2 protein in 10% glycerol was loaded slowly onto the column. The column was then washed with five CV of buffer A-1 (20 mM Tris-HCl pH 7.5, 100 mM $NH_4Cl$, 1 mM $MgCl_2$, 0.5 mM EDTA, 2 mM dithiothreitol), and crude ribosomes in buffer A-1 were loaded onto the column. The column was washed with five CV of buffer A-1, 10 CV of buffer A-250 (buffer A-1 with 250 mM $NH_4Cl$), and then tagged ribosomal subunits were eluted with a 10 CV gradient of 0–10 mM maltose in buffer A-1. The purified 50S subunits were concentrated using 100 kDa cut off spin filters (Millipore), quantified, and flash frozen in liquid nitrogen. The purity of the resulting MS2-tagged 50S subunits was verified to be 98% using PCR primers (Table S3) spanning the MS2-binding RNA hairpin sequence as previously described[1]. WT untagged 30S and 50S ribosomal subunits were purified as previously described[1].

*Quantification of activity using a bioluminescence complementation assay.*

A HiBit peptide (peptide-1, Table S4) was generated using *in vitro* transcription/translation (IVTT) from a template containing a T7 promoter, a ribosome binding site, and the coding sequence for the HiBit sequence[3]. DNA templates were obtained as Ultramers from IDT (Table S3) and PCR amplified (5 ng of template, 0.5 µM forward primer, 0.5 µM reverse primer and Taq 2X Master Mix). Templates were purified using a DNA Clean and Concentrator Kit (Zymo Research). IVTT reactions were performed using a Δribosome Purexpress kit (NEB) where ribosomes were omitted from the reactions and purified, tagged WT or mutant ribosomes were used. All Purexpress reagents were added on ice. Then 0.5 µM of purified 30S subunits, 0.25 µM of either WT or mutant ribosomes, 0.1 µM of LgBiT, and a 1:50 dilution of Nano-Glo substrate (Promega) were added. Finally, the reaction was initiated by addition of 100 ng of DNA template, placed in a 384-well plate, and luminescence was measured over time at 37 ºC in a Spark Plate Reader (Tecan).

*In vitro translation of peptides containing a variable number of prolines.*

Peptides-2-4 (Table S4) were generated using *in vitro* transcription/translation (IVTT) from a template containing a T7 promoter, a ribosome binding site, the coding sequence for a short peptide with a FLAG-tag, and a variable number of prolines. DNA templates were obtained as Ultramers (Table S3) from IDT and PCR amplified (5 ng of template, 0.5 µM forward primer, 0.5 µM reverse primer and Taq 2X Master Mix). Templates were purified using a DNA Clean and Concentrator Kit (Zymo Research). IVTT reactions were performed using a Δribosome Purexpress kit (NEB) where ribosomes were omitted from the reactions. All Purexpress reagents were added on ice, then 1 µM of purified 30S subunits, 0.5 µM of either WT or mutant 50S subunits. Finally, the reaction was initiated by addition of 100 ng of DNA template. Reactions were incubated at 37 ºC for 4 hours. In experiments testing the effect of EF-P, 5 µM EF-P (Cosmo Bio) was included in the IVTT reactions.

#### *Purification and analysis of FLAG-tagged peptides*

Anti-FLAG M2 Magnetic Beads (Millipore Sigma) were used to purify IVTT generated, FLAG-tagged peptides for downstream analysis. Anti-FLAG beads (12.5 µL) were washed with 100 µL wash buffer (50 mM Tris HCl pH 7.5, 150 mM NaCl) and the supernatant was removed from the beads. IVTT reactions were applied to the anti-FLAG beads and incubated at room temperature for 30 minutes. The supernatant was then removed, and the beads were washed 2X with 100 µL of wash buffer, and once with 100 µL of RNase free water. Elution buffer (12.5 µL) (100 mM glycine pH 2.8) was added to the beads and incubated at room temperature for 10 min[4]. The supernatant was removed and analyzed by LCMS. Samples were resolved on an an Agilent 1290 Infinity II HPLC using an Eclipse XDB C18 Column (1.8 µm, 2.1mm x 50 mm, 25 ºC) (Agilent) with mobile phases A (water with 0.1 % [v/v] formic acid) and B (acetonitrile with 0.1 % [v/v] formic acid). The method began with a linear gradient from 0% B to 1% B for one minute, then a linear gradient was used from 1-91% B over five minutes at a flow rate of 0.5 mL/min, then subsequently washed with 91% B for 0.6 minutes, then a linear gradient decreased B from 91% to 1% for 1.1 minutes, finally the column was equilibrated 1% B for 1.3 minutes. The mass spectra of each sample was obtained on an Agilent 6530B QTOF AJS-ESI. The masses were analyzed using the MassHunter software (Agilent) by extracting the major ion corresponding to each peptide species (± 100 ppm) and integrating the subsequent extracted ion chromatogram[4].

#### *Transcription of tRNA$^{Pyl(Ser)}$*

A DNA template for *E. coli* tRNA$^{Pyl(Ser)}$ was obtained as an Ultramer from IDT (Table S3). The template was PCR amplified with forward and reverse primers corresponding to the template (Table S3). Reverse primers included a 2′-O-methyl modification to reduce heterogeneity on the 3′-end of the sequence during transcription[5]. PCR reactions were performed with 5 ng of template, 0.5 µM forward primer, 0.5 µM reverse primer and Q5 High-Fidelity 2X Master Mix (New England BioLabs). *In vitro* transcription of tRNAs was carried out in transcription buffer (50 mM Tris pH 7.5, 2 mM spermidine, 0.1% triton X100), 12 mM $MgCl_2$, 8 mM DTT, 2 mM each nucleotide (NEB), 1 U/µL of murine RNase inhibitor (NEB), 0.5 U/µL *E. coli* inorganic pyrophosphatase (NEB), 2 U/µL of T7 RNA polymerase (NEB) and 15% v/v of the PCR reaction containing the amplified DNA template. The reaction was incubated at 37 ºC for 4 hours, after which the products were precipitated with EtOH and purified on a 10% polyacrylamide, 7 M urea, 0.5X TBE (50 mM Tris, 50 mM boric acid, 1 mM EDTA) gel. Bands corresponding to tRNA were excised from the gel and the RNA was eluted for 2 hours in 300 mM KCl after which the gel pieces were removed and the tRNA was precipitated with EtOH. The pellets were resuspended in water and stored in a -80 ºC freezer prior to use.

#### *Enzymatic acylation of tRNA$^{Pyl(Ser)}$ with either α-BocK or (R)-β$^2$-OH-BocK*

tRNA$^{Pyl(Ser)}$ was enzymatically acylated with α-BocK or *(R)*-*β*$^2$-OH-BocK using PylRS. tRNA$^{Pyl(Ser)}$ was incubated with 12.5 µM PylRS in aminoacylation buffer (100 mM HEPES pH 7.5, 10 mM $MgCl_2$, 4 mM DTT, and 10 mM ATP) along with 0.004 U/µL inorganic pyrophosphatase and 5 mM of α-BocK or *(R)*-*β*$^2$-OH-BocK. The reactions were incubated at 37 ºC for 2 hrs and purified using a Zymo RNA Clean and Concentrate kit. Acylation was confirmed and quantified by intact tRNA mass spectrometry (Majumdar et al. 2026)

#### *Flexizyme acylation of tRNA$^{Pyl(Ser)}$* with either *(R)-β$^2$-NH$_2$-Phe or (S)-β$^2$-NH$_2$-Phe*

A flexizyme (eFx) was used to promote the reaction between unacylated tRNA$^{Pyl(Ser)}$ and *(R)-β2*-NH2-Phe or *(S)-β2*-NH2-Phe preactivated as cyanomethyl esters (*(R)-β2*-PheCME and *(S)-β2*-PheCME respectively). The tRNA (25 μM) was added to a mixture containing 25 μM eFx and 50 mM HEPES pH 9.0 and then co-folded. Following co-folding, the following reagents were added in the subsequent order; 600 mM $MgCl_2$, 5 mM *(R)-β2*-PheCME or *(S)-β2*-PheCME monomer dissolved in DMSO, and an additional 10% DMSO supplement to aid in monomer solubility. The mixture was incubated at 4 ºC and allowed to react for 2 hours. The charged tRNA was purified by first quenching with 3 M NaOAc (pH 5.2) before 4 equivalents of 100% molecular biology grade ethanol was added and the tRNA was allowed to precipitate at -80 ºC for 1 hour. The precipitated tRNA was then quickly isolated by rapidly pelleting at 21,000 xG for 15 minutes in a precooled benchtop centrifuge. The supernatant was then carefully removed, the pellet was washed with an ice-cold mixture of 70% EtOH and 0.1 mM NaOAc (pH 5.2) which was also removed following a 5 minute centrifugation at 21,000 xG. The pellet was then redissolved in RNAse-free water and analyzed by intact tRNA mass spectrometry. Percent yields were calculated by extracting the major ion corresponding to each tRNA species and integrating the subsequent extracted ion chromatogram

*In vitro translation of peptides containing either α-BocK or $β^2$-OH-BocK.*

Peptides-5-6 (Table S4) were generated using *in vitro* transcription/translation (IVTT) from a template containing a T7 promoter, a ribosome binding site, the coding sequence for a short peptide with a FLAG-tag, and a variable number of serine codons. The serine codons were decoded by either 3′-$tRNA^{Pyl}$-α-BocK or 3′-$tRNA^{Pyl}$-*(R)*-$β^2$-OH-BocK. DNA templates were obtained as Ultramers (Table S3) from IDT and PCR amplified (5 ng of template, 0.5 μM forward primer, 0.5 μM reverse primer and Taq 2X Master Mix). Templates were purified using a DNA Clean and Concentrator Kit (Zymo Research). IVTT reactions were performed using a combination of a Δribosome Purexpress kit (NEB) and Δamino acid/tRNA Purexpress kit (NEB) where ribosomes and amino acids were removed from the reaction. All Purexpress reagents were added on ice, then 1 μM of purified 30S subunits, 0.5 μM of either WT or mutant ribosomes, and an amino acid mixture without serine. Finally, the reaction was initiated by addition of 100 ng of DNA template. Reactions were incubated at 37 ºC for 4 hours.

*In vitro translation of peptide to test tryptophan codon stalling strategy.*

Peptide-7 (Table S4) was generated using *in vitro* transcription/translation (IVTT) from a template containing a T7 promoter, a ribosome binding site, the coding sequence for a short peptide with a FLAG-tag, and a variable number of prolines followed by a tryptophan codon. DNA templates were obtained as Ultramers (Table S3) from IDT and PCR amplified (5 ng of template, 0.5 μM forward primer, 0.5 μM reverse primer and Taq 2X Master Mix). Templates were purified using a DNA Clean and Concentrator Kit (Zymo Research). IVTT reactions were performed using a combination of a Δribosome Purexpress kit (NEB) and Δamino acid/tRNA Purexpress kit (NEB) where ribosomes and amino acids were removed from the reaction. All Purexpress reagents were added on ice, then 1 μM of purified 30S subunits, 0.5 μM of either WT or mutant ribosomes, and an amino acid mixture with or without tryptophan. Finally, the reaction was initiated by addition of 100 ng of DNA template. Reactions were incubated at 37 ºC for 4 hours. FLAG-tagged peptides were purified and analyzed using LCMS as described above.

*Cryo-EM complex preparation.*

A longer stalling sequence was used to isolate ribosomes for cryo-EM (Table S3). IVTT of the sequence containing a T7 promoter, a ribosome binding site, the coding sequence for an N-terminal FLAG-tag, a longer peptide sequence, three prolines followed by a tryptophan codon. DNA templates were obtained as a G-Block (Table S3) from IDT and PCR amplified (5 ng of template, 0.5 μM forward primer, 0.5 μM reverse primer and Taq 2X Master Mix). Templates were purified using a DNA Clean and Concentrator Kit (Zymo Research). IVTT reactions were performed using a combination of a Δribosome Purexpress kit (NEB) and Δamino acid/tRNA Purexpress kit (NEB) where ribosomes and amino acids were removed from the reaction. All Purexpress reagents were added on ice, then 2 μM of purified 30S subunits, 1.5 μM of either

WT or mutant ribosomes, and an amino acid mix without tryptophan. Finally, the reaction was initiated by addition of 100 ng of DNA template. Reactions were incubated at 37 ºC for 5 hours.
Ribosome complexes were purified from the solution using Anti-FLAG M2 Magnetic Beads (Millipore Sigma). Anti-FLAG beads (50 µL) were washed with 750 µL of buffer (20 mM HEPES HCl pH 7.5, 50 mM $NH_4Cl$, 50 mM KCl, 15 mM $MgCl_2$) and the supernatant was removed from the beads. IVTT reactions were applied to the anti-FLAG beads and incubated at room temperature for 20 minutes with mixing. The supernatant was then removed, and the beads were washed 2X with 500 µL of buffer. Elution was done with 50 µL of 100 µM 3x FLAG-peptide in the buffer and incubated at room temperature for 20 minutes. The supernatant was collected and applied to a 100 kDa cutoff spin filter and subsequently washed and concentrated with buffer. The complexes were frozen in liquid nitrogen and thawed just prior to plunge freezing on grids.

*Cryo-EM sample preparation.*
The samples were prepared for imaging on 300 mesh R1.2/1.3 UltrAuFoil grids (Quantifoil) with an additional layer of amorphous carbon support film. Before applying the sample, grids were glow discharged in a PELCO easiGlow at 0.37 mBar and 20 mAmp for 12 seconds. 4 µL of sample was deposited onto each grid and incubated for 1 minute. Grids were blotted and plunge-frozen in liquid ethane with an FEI Mark IV Vitrobot using the following settings: 4 °C, 100% humidity, blot force 3, and blot time 2. Grids were clipped for autoloading and stored in liquid nitrogen.

*Cryo-EM data collection.*
Cryo-EM data collection was conducted after screening grids for ribosome density and grid integrity. Dose fractionated movies were collected on a Titan Krios G3i microscope at an accelerating voltage of 300 kV and with a BIO Quantum energy filter. Movies were recorded on a GATAN K3 direct electron detector operated in CDS mode. A total dose of 40 $e^-/Å^2$ was split over 40 frames per movie. The physical pixel size was set to 0.81 Å and the super-resolution pixel size to 0.405 Å. Data collection was automated with SerialEM[6], which was also used for astigmatism correction by CTF and coma-free alignment by CTF. The defocus ramp was set to range between -0.5 and -1.5 µm.

*Cryo-EM image processing.*
Data processing was performed using cryoSPARC v4.6.2[7]. Patch motion correction was used to motion correct the imported movies and CTF Find[8] was used to estimate CTF parameters and micrographs with poorly fitting CTF estimates were omitted. Particles were picked using Blob Picker and extracted at one-fourth binning of the full box size and subjected to 2D Classification to select 70S particles. 3D classification was performed using Heterogeneous Refinement using a 70S reference volume generated from the coordinates of 7K00 using EMAN2[9]. Classes that corresponded to a 70S were then selected and subjected to 3D Classification on P-site tRNA. Classes that appeared to have a P-site tRNA and an empty A-site. Resulting particles were then extracted at the full box size and subjected to Homogeneous Refinement. A 3D Classification was performed on the exit tunnel peptide and classes that had a peptide present in the exit tunnel were subjected to another Homogeneous Refinement. The global resolution of the final maps were 1.88 Å for WT complexes and 2.49 Å for mutant A2062U complexes.

*Modeling.*

PDB entry 8QOA containing the stalled peptide SecM in the exit tunnel of the ribosome was used as the starting model[10]. The A-site tRNA was removed and the P-site tRNA was remodeled to $tRNA^{Pro}$. The appropriate mRNA sequence was added, and the five amino acids connected to the P-site tRNA were added. The remainder of the nascent peptide was removed. Real space refinement of the coordinates was performed in PHENIX[11] and further adjustments to the model were done manually in Coot[12] in the vicinity of the PTC. Additions and deletions to the model included $Mg^{2+}$ and $K^{+}$ ions and water molecules in the vicinity of the PTC.

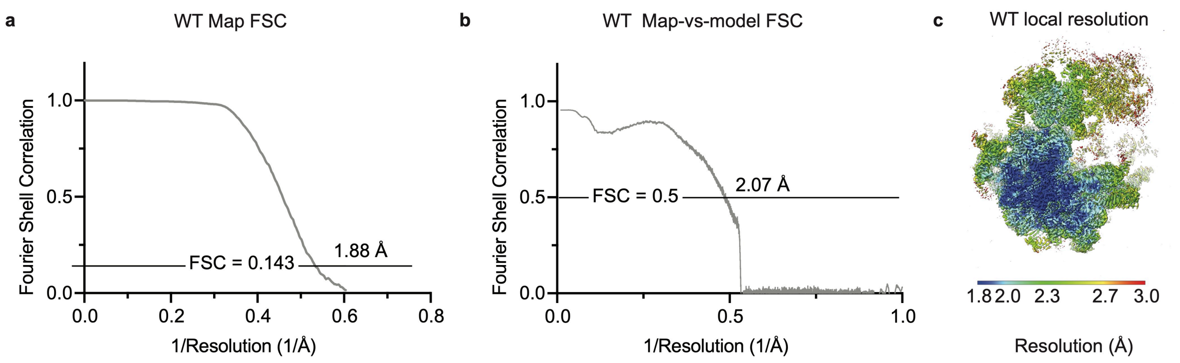


**Figure S1:** Resolution estimates 70S WT ribosome complexed with the poly-Pro stalling sequence. **a,** FSC curve of the 70S map (gray) showing the global resolution of the map is 1.88 Å at the gold-standard cutoff value of 0.143. **b,** Map-vs-model resolution is at 2.07 Å at FSC cutoff value of 0.5. **c,** 70S map color coded by local resolution values.

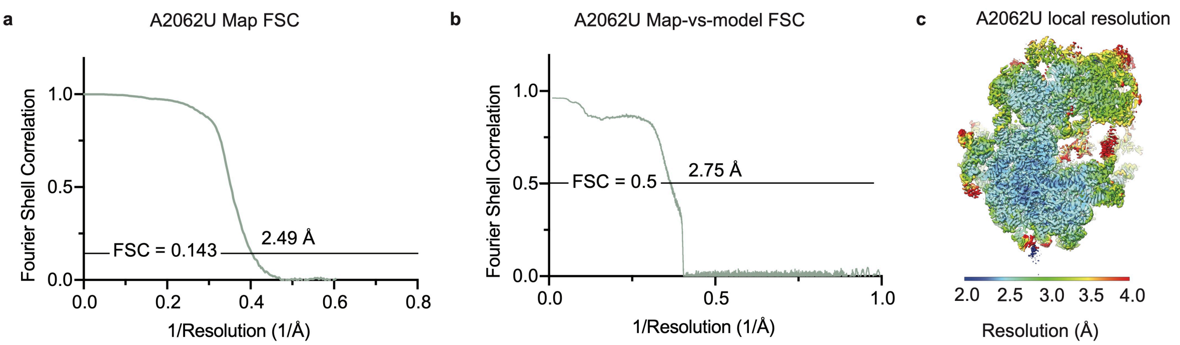


**Figure S2:** Resolution estimates 70S A2062U mutant ribosome complexed with the poly-Pro stalling sequence. **a,** FSC curve of the 70S map (green) showing the global resolution of the map is 2.49 Å at the gold-standard cutoff value of 0.143. **b,** Map-vs-model resolution is at 2.75 Å at FSC cutoff value of 0.5. **c,** 70S map color coded by local resolution values.

Data processing steps for WT ribosome P-site tRNA$^{Pro}$-3-Pro-peptide

Data collection
9,824 movies

↓

Motion correction
Select micrographs
with good CTF estimates
9,639 micrographs

↓

Autopicking 70S particles
1,119,091 particles

↓

2D class
1,090,656 particles

↓

3D classification
Select 70S classes
743,095 particles

↓

Masked classification
Select particles with
occupied P site
659,879 particles

↓

Masked classification
Select particles with
occupied peptide
227,702 particles

↓

3D Refinement
1.88 Å

**Figure S3:** Cryo-EM data processing workflow for the WT 70S ribosome with a polyproline nascent chain.

Data processing steps for A2062U mutant ribosome P-site $tRNA^{Pro}$-3-Pro-peptide

Data collection
8,242 movies

↓

Motion correction
Select micrographs
with good CTF estimates
7,728 micrographs

↓

Autopicking 70S particles
1,058,420 particles

↓

2D class
1,058,420 particles

↓

3D classification
Select 70S classes
552,986 particles

↓

Masked classification
Select particles with
occupied P site
210,657 particles

↓

Masked classification
Select particles with
unoccupied A site
115,708 particles

↓

Masked classification
Select particles with
occupied peptide
39,287 particles

↓

3D Refinement
2.49 Å

**Figure S4:** Cryo-EM data processing workflow for the mutant A2062U 70S ribosome with a polyproline nascent chain.

**Table S1:** Data collection and processing for the WT 70S ribosome and the mutant A2062U ribosome with a poly-proline nascent chain.

| **Complex** | ***WT*** | ***A2026U*** |
|---|---|---|
| Magnification | 104,414 | 104,452 |
| Voltage (kV) | 300 | 300 |
| Electron Exposure ($e^-$/Å$^2$) | 40 | 40 |
| Defocus Range (μm) | -0.5/-1.5 | -0.5/-1.5 |
| Pixel Size (Å) | 0.8246 | 0.8243 |
| Symmetry Imposed | C1 | C1 |
| Initial Particle Images | 1,119,091 | 1,058,420 |
| Final Particle Images | 227,702 | 39,287 |
| Map Resolution (Å) | 1.88 | 2.49 |
| FSC Threshold | 0.143 | 0.143 |

**Table S2:** Model refinement statistics for the WT 70S ribosome and the mutant A2062U ribosome with a poly-proline nascent chain.

| **Model component** | ***WT*** | ***A2062U*** |
|---|---|---|
| Model resolution (Å) | 2.0 | 2.7 |
| FSC threshold | 0.5 | 0.5 |
| Map sharpening *B* factor ($Å^2$) | 32 | 40.8 |
| Model composition | | |
| Non-hydrogen atoms | 145,489 | 142,167 |
| $Mg^{2+}$ ions | 278 | 240 |
| Waters | 5482 | 2176 |
| Mean *B* Factors ($Å^2$) | | |
| RNA | 68.66 | 68.97 |
| Protein | 78.88 | 72.92 |
| Waters | 48.34 | 56.50 |
| Other | 66.33 | 78.64 |
| R.m.s. deviations from ideal values | | |
| Bond (Å) | 0.005 | 0.005 |
| Angle (°) | 0.502 | 0.589 |
| Molprobity score | 1.69 | 1.94 |
| Clash Score | 7.42 | 8.41 |
| Rotamer outliers (%) | 1.89 | 2.91 |
| Ramachandran plot | | |
| Favored (%) | 97.71 | 97.29 |
| Allowed (%) | 2.24 | 2.69 |
| Outliers (%) | 0.05 | 0.02 |
| RNA validation | | |
| Angles outliers (%) | 0.002 | 0.001 |
| Sugar pucker outliers (%) | 0.270 | 0.315 |
| Average suiteness | 0.603 | 0.588 |

**Table S3:** DNA sequences.

| Description | Sequence |
|---|---|
| A2062U Cloning Fwd. | AGACGGAAAGTCCCCGTGAACCTTTAC |
| A2062U Cloning Rev. | TGCCGCGGGTACACTGC |
| Templates Primer Fwd. | GCGAATTAATACGACTCACTATAGGGTTAACTTTAAC |
| Templates Primer Rev. | TGACTAAACCCCTCCGTTTAGAGAGG |
| MS2 Primer Fwd. | CTTGCCCCGAGATGAGTTCTCCC |
| MS2 Primer Rev. | GTACCGGTTAGCTCAACGCATCGCT |
| HiBit Peptide Template | GCGAATTAATACGACTCACTATAGGGTTAACTTTAACAAGGAGAAAAACATGGT GAGCGGCTGGCGCCTGTTTAAAAAAATTAGCTAACTAGCATAACCCCTCTCTA AACGGAGGGGTTTAGTCA |
| 2-Proline Template | GCGAATTAATACGACTCACTATAGGGTTAACTTTAACAAGGAGAAAAACATGTA CAAGTACAAAAAGTACAAACCGCCGGGTGACTACAAAGACGATGACGACAAGT AACTAGCATAACCCTTCTCTAAACGGAGGGGTTTAGTCA |
| 3-Proline Template | GCGAATTAATACGACTCACTATAGGGTTAACTTTAACAAGGAGAAAAACATGTA CAAGTACAAAAAGTACAAACCGCCGCCGGGTGACTACAAAGACGATGACGAC AAGTAACTAGCATAACCCTTCTCTAAACGGAGGGGTTTAGTCA |
| 4-Proline Template | GCGAATTAATACGACTCACTATAGGGTTAACTTTAACAAGGAGAAAAACATGTA CAAGTACAAAAAGTACAAACCGCCGCCGCCGGGTGACTACAAAGACGATGAC GACAAGTAACTAGCATAACCCTTCTCTAAACGGAGGGGTTTAGTCA |
| 2-Serine Template | GCGAATTAATACGACTCACTATAGGGTTAACTTTAACAAGGAGAAAACATGGCT AGCAGCAAAGACTACAAAGACGATGACGACAAGTAACTAGCATAACCCCTCTC TAAACGGAGGGGTTTAGTCA |
| 3-Serine Template | GCGAATTAATACGACTCACTATAGGGTTAACTTTAACAAGGAGAAAACATGGCT AGCAGCAGCAAAGACTACAAAGACGATGACGACAAGTAACTAGCATAACCCCT CTCTAAACGGAGGGGTTTAGTCA |
| Trp-Stall for LCMS Template | GCGAATTAATACGACTCACTATAGGGTTAACTTTAACAAGGAGAAAAACATGTA CAAGTACAAAAAGTACAAACCGCCGCCGTGGGGTGGTGACTACAAAGACGAT GACGACAAGTAACTAGCATAACCCTTCTCTAAACGGAGGGGTTTAGTCA |
| tRNA-Pyl-Ser Primer Fwd. | AATTCCTGCAGTAATACGACTCA |
| tRNA-Pyl-Ser Primer Rev. | TmGGCGAGAGACCGGG |
| tRNA-Pyl-Ser-Template | ATTCCTGCAGTAATACGACTCACTATAGGGGGACGGTCCGGCGACCAGCGGG TCTGCTAAACCTAGCCAGCGGGGTTCGACGCCCCGGTCTCTCGCCA |

| | |
|---|---|
| Trp-Stall for Cryo-EM Template | GCGAATTAATACGACTCACTATAGGGTTAACTTTAACAAGGAGAAAAACATGGGCAGCAGCCATCATCATCATCATCACGATTACAAGGATGACGACGATAAGGCTAGCAGCAGCGGTACCGGCAGCGGCGAAAACCTCTATTTTCAGGGTAGTGCGCAAGCATGCGTGAGTGGAATACTGACGCGCCGACAGTTTGGTAAACGCTACTTCCCGCATCTCTTATTAGGGATGGTTGCGGCGAGTTTAGGTTTGCCTGCGCTCAGCTACAAGAAGTACAAAAAGTACAAACCGCCGCCGTGGGGTGGTTTAGGTTTGCCTGCGCTCAGCTAACTAGCATAACCCCTCTCTAAACGGAGGGGTTTAGTCA |

**Table S4:** Peptide sequences.

| Label | Description | Sequence |
|---|---|---|
| Peptide-1 | HiBit | MVSGWRLFKKIS |
| Peptide-2 | 2-Proline | MYKKYKKYKPPGDYKDDDDK |
| Peptide-3 | 3-Proline | MYKKYKKYKPPPGDYKDDDDK |
| Peptide-4 | 4-Proline | MYKKYKKYKPPPPGDYKDDDDK |
| Peptide-5 | 1-Serine | MASKDYKDDDDK |
| Peptide-6 | 2-Serine | MASSKDYKDDDDK |
| Peptide-7 | 3-Serine | MASSSKDYKDDDDK |
| Peptide-8 | Trp-Stall | MYKKYKKYKPPPWGDYKDDDDK |
